# Prolonged non-suppressible viremia sustained by a clonally expanded, genomically defective provirus with an immune-evasive HIV protein expression profile

**DOI:** 10.1101/2025.05.16.654538

**Authors:** F. Harrison Omondi, Yurou Sang, Francis Mwimanzi, Peter K. Cheung, Zerufael Derza, Winnie Dong, Evan Barad, Kieran Anderson, Vitaliy Mysak, Viviane D. Lima, Mark Hull, Chanson J. Brumme, Marianne Harris, Julio S.G. Montaner, Silvia Guillemi, Mark A. Brockman, Zabrina L. Brumme

## Abstract

Antiretroviral therapy (ART) potently inhibits HIV replication but does not eliminate HIV proviruses integrated within the genomes of infected cells. Though ART normally suppresses HIV to clinically undetectable levels in blood, some individuals experience non-suppressible viremia (NSV) that is not attributable to suboptimal drug exposure (*e.g.* due to incomplete medication adherence or pharmacological issues) nor emerging HIV drug resistance, and that does not resolve following regimen modification. We now understand that NSV can originate from expanded cell clones harboring genetically identical proviruses that reactivate *en masse* to produce clinically detectable viremia. NSV can even originate from proviruses with genomic defects, particularly in HIV’s major splice donor (MSD) site, that render the viremia non-infectious. But, because only a limited number of such cases have been described, all in HIV subtype B, the mechanisms that allow cells harboring such proviruses to produce prolonged viremia without being eliminated remain incompletely understood. We characterized a case of MSD-defective, replication-impaired NSV that lasted >4 years in an individual with non-B HIV. Our results reveal that proviruses with MSD deletions can persist by integrating into minimally differentiated CD4+ T-cell subsets that then give rise to the full spectrum of memory and effector subpopulations, and by exhibiting an HIV protein expression profile that would allow these cells to evade cellular and humoral immunity. Our results highlight the need to better understand the biological implications of persistent HIV protein and virion production by genomically defective, clonally expanded proviruses, and for clinical guidelines to acknowledge this type of viremia.

**Importance:** HIV cure efforts focus on eliminating the viral reservoir, strictly defined as cells harboring proviruses that can produce infectious HIV. But this definition excludes proviruses with defects in HIV’s major splice donor site (MSD), which persist readily. Our results confirm that clonally expanded, MSD-defective proviruses can produce HIV transcripts, proteins and clinically-detectable viremia over long periods, underscoring the need to investigate the potential long-term biological consequences of these activities. Our findings also have clinical implications. Most HIV treatment guidelines don’t acknowledge that persistent viremia during ART can originate from clonally-expanded proviruses, and all recommend that viremia >200 HIV RNA copies/mL be managed as virologic failure. Clear clinical frameworks should be developed to discriminate persistent viremia that is due to drug adherence, pharmacokinetics or emerging drug resistance (that is clinically actionable), from NSV (that is not). Genotyping of HIV’s MSD could also help assess potential transmission risk during NSV.

## INTRODUCTION

Antiretroviral therapy (ART) halts Human Immunodeficiency Virus (HIV) replication and normally suppresses the virus to clinically undetectable levels in blood. Despite this, some individuals receiving ART experience sustained low-level viremia with no obvious cause (1–5). Such cases raise concerns about incomplete ART adherence, suboptimal drug exposure due to pharmacological issues, emerging drug resistance and possible virologic failure (6–9), but this phenomenon also occurs in highly adherent individuals and in the absence of new drug resistance mutations (1, 2, 5, 10). In such cases, the viremia is typically not resolvable through ART modification or intensification, hence the term “non-suppressible viremia” (NSV) (4, 11).

HIV, like all retroviruses, persists as an integrated provirus within infected cells, and these “reservoir cells” can clonally expand and reactivate at any time to produce infectious HIV, provided the provirus is genomically intact (12–16). It is for this reason that ART needs to be taken for life. Recently however, it has been demonstrated that NSV can originate from expanded cell clones that harbor genetically identical HIV proviruses that reactivate *en masse* to release sufficient virus detectable by conventional assays (3, 4, 17, 18). While ART blocks these viruses from infecting new cells, it neither halts clonal expansion nor eliminates the cells themselves, allowing HIV to be released over long periods.

Notably, NSV is not always infectious. It can sometimes originate from proviruses with defects in the 5’ leader region of the HIV genome, specifically within the Major Splice Donor (MSD) site (HXB2 nucleotide 743) (3, 17), which is used to produce the full complement of spliced viral mRNA transcripts (19–21). Though proviruses with MSD mutations/deletions can produce some viral transcripts, proteins, and even viral particles (3, 12, 22), MSD defects generally alter transcript composition, reduce viral RNA packaging efficiency, and impair virion infectivity (22, 23). Nevertheless, the full impacts of MSD defects on HIV spliced transcript production and protein expression remain incompletely understood, as do the mechanisms that allow clonally expanded cells harboring MSD-defective proviruses to produce viremia for years without being eliminated by the immune system. This is in part because, to our knowledge, only eight such cases have been described to date (3, 17), all of which were HIV subtype B infections where most NSV clones had sizeable (>20 base) MSD deletions.

To better understand this intriguing phenomenon, we longitudinally characterized a case of replication-impaired NSV in an individual with a non-subtype B infection that lasted for >4 years and substantially exceeded the >200 HIV RNA copies/mL threshold that commonly defines virologic failure (6, 7, 24). The viremia in this case was defective due to a novel three base MSD deletion. We identified the CD4+ T-cell subsets harboring the NSV-causing provirus in blood, and characterized intracellular HIV transcription and protein expression profiles to examine how these cells evaded immune detection. Our results reveal that proviruses with MSD deletions can persist by initially integrating into minimally differentiated CD4+ T-cell subsets and by exhibiting an HIV protein expression profile that facilitates immune evasion. Our study highlights the need to better understand the potential biological implications of long-term virus release by clonally expanded proviruses, and for HIV treatment guidelines to acknowledge that persistent viremia >200 HIV RNA copies/mL does not always constitute virologic failure.

## RESULTS

### Participant characteristics and sampling

The participant, a Black male in his 50s, was diagnosed with HIV in 2003, though he believes that he acquired HIV around 1990 (**Figure 1A**). By Fall 2006, his CD4+ T-cell count had reached a nadir of 220 cells/mm^3^, his plasma viral load exceeded 100,000 HIV RNA copies/mL, and he initiated ART comprising tenofovir, emtricitabine and efavirenz. This regimen was maintained until May 2018, when efavirenz was replaced with the integrase strand transfer inhibitor (INSTI) elvitegravir boosted with cobicistat. With the exception of isolated viremia “blips”, plasma viral loads remained suppressed on ART until late 2019, when a viral load of 43 copies/mL was recorded. In early 2020, ART was changed to emtricitabine, tenofovir alafenamide and the INSTI bictegravir, to simplify to a one-pill-per-day regimen. The viral load measurement one month later was 83 copies/mL. A >4-year period of persistent low-level viremia then followed, reaching up to 709 copies/mL, that did not resolve despite regimen modification, including intensification to 4 active drug classes. No adherence concerns were noted, the presence of antiretrovirals in plasma was confirmed using a validated mass spectrometry assay (25, 26), and clinical HIV drug resistance genotyping performed 11 times during the viremic period revealed a single unchanging HIV sequence with no drug resistance mutations that was distinct from prior genotypes (**Figure 1B**). These observations indicated that the viremia was not due to adherence nor pharmacokinetic issues (9) nor *de novo* HIV drug resistance (8), but likely due to HIV release from a clonally expanded provirus (3, 4, 17). We henceforth refer to this as “non-suppressible viremia” (NSV).

**Figure 1.**
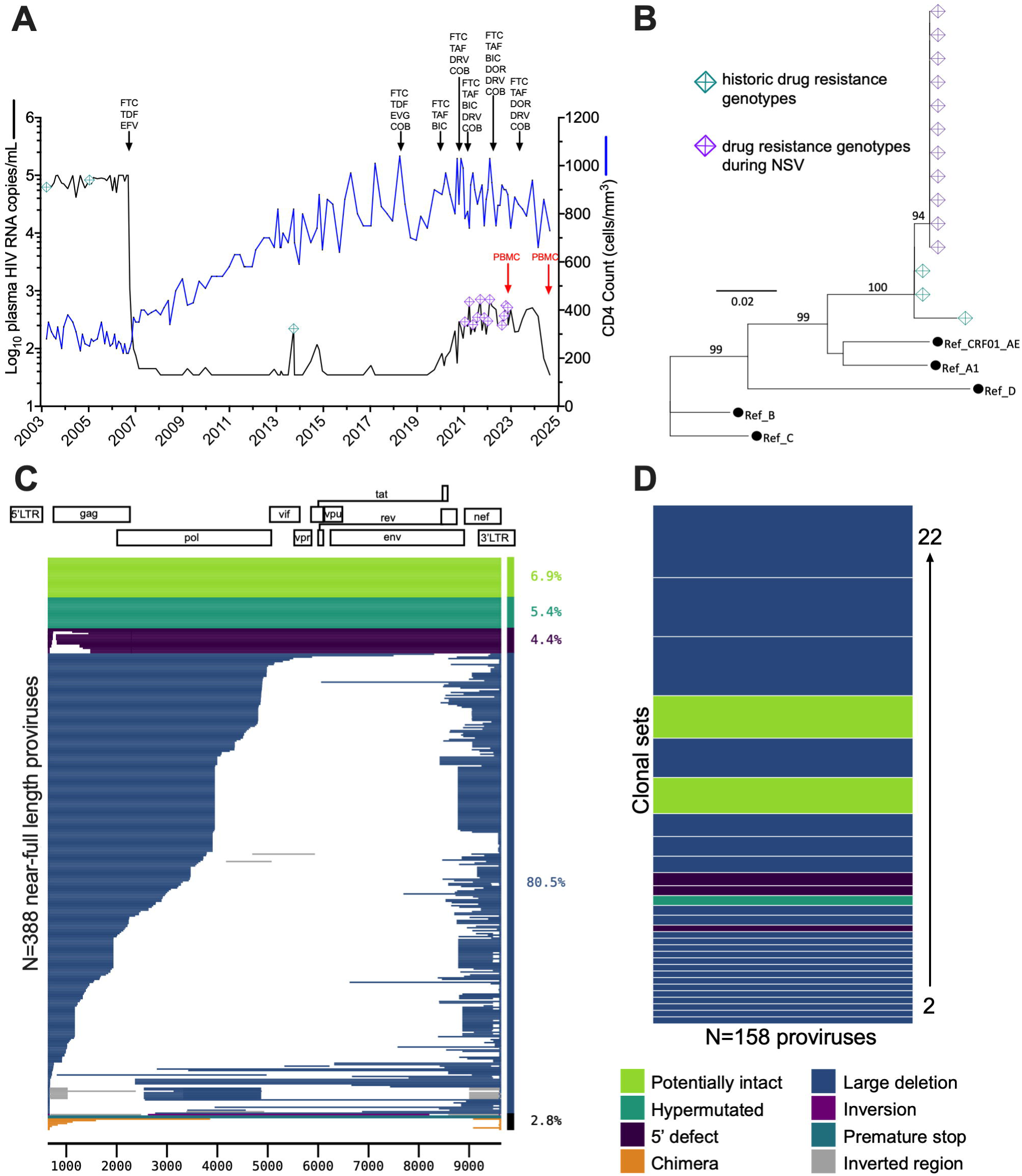
Participant clinical history and sampling timeline. (A) Plasma viral load (black line), CD4 count (blue line) and antiretroviral history. FTC = emtricitabine; TDF = tenofovir disoproxil fumarate; EFV = efavirenz; EVG = elvitegravir; COB = cobicistat; TAF = tenofovir alafenamide; BIC = bictegravir; DRV = darunavir; DOR = doravirine. NSV occurred from approximately Jan 2020-Sept 2024. Teal and purple crossed diamonds denote HIV drug resistance testing performed before and during NSV, respectively. Red arrows denote peripheral blood mononuclear cells (PBMC) sampling. (B) Maximum likelihood within-host phylogeny of partial *pol* sequences from drug resistance genotypes performed before and during NSV. Black circles denote reference sequences. Numbers indicate branch support values. Scale is in estimated substitutions per nucleotide site. (C) Proviral landscape plot depicting 388 near-full-length proviruses isolated from blood CD4+ T-cells during NSV, colored according to genomic integrity (white denotes deletions). The frequencies of each provirus type are shown on the right of the plot. (D) Summary of proviral clones with 100% sequence identity, colored according to genomic integrity. Bar height corresponds to clone size. The fourth-largest clone, in light green, is the one that matches the NSV.

### The NSV arises from a clonally expanded provirus

Approximately 3 years after the NSV began, the participant provided blood from which we isolated 388 near-full-length proviruses from CD4+ T-cells by single-genome amplification (**Figure 1C**). This revealed the virus was a complex recombinant of subtypes A1, G, CRF01-AE, and CRF02-AG (**Figure S1**). Of the proviruses recovered, 6.9% were bioinformatically classified as potentially genetically intact, while 4.4% harbored obvious defects in the 5’ leader region. Consistent with clonal expansion, we observed numerous identical proviruses, including two sizeable clones that were potentially intact (**Figure 1D**, green). The *pol* region of the larger of these clones exactly matched the longitudinal genotypes that had been performed during the NSV (**Figure 2A**). Partial 5’ leader and *env* regions single-genome amplified from plasma HIV RNA during NSV also exactly matched this same proviral clone (**Figure S2**). This indicated that the NSV originated from a clonally expanded CD4+ T-cell clone that was present in blood.

**Figure 2.**
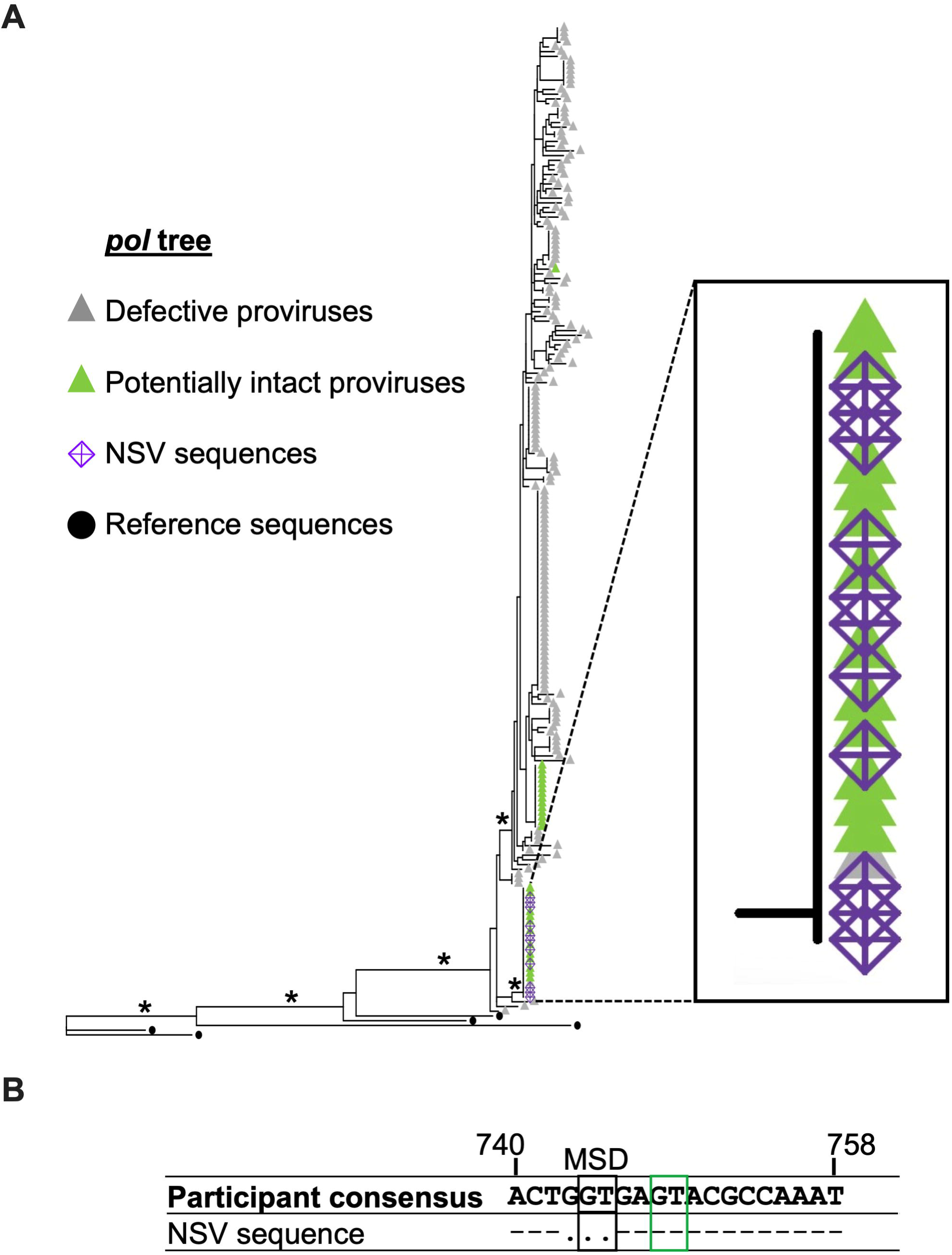
An exact match was found between NSV and a clonally expanded provirus. (A) Maximum likelihood within-host phylogeny inferred from 198 non-hypermutated proviral sequences that contained the partial *pol* region genotyped for HIV drug resistance testing (where green and grey triangles indicate potentially intact and defective proviruses, respectively), along with the 11 identical plasma HIV RNA genotypes performed during the NSV (purple crossed diamonds). The inset shows the proviruses identical to the NSV sequences. Black filled circles denote reference sequences. Asterisks denote branch support values >80%. Scale is in estimated substitutions per nucleotide site. (B) Nucleotide alignment of the participant’s proviral consensus sequence versus the NSV sequence in the major splice donor (MSD) region, with HXB2 genomic coordinates shown above. The black box denotes the MSD, while the green box denotes a known cryptic splice donor site.

### The NSV provirus produced replication-impaired virions due to a three-base MSD deletion

Though the NSV provirus was initially classified as potentially intact, it harbored a three-base “GGT” deletion in HIV’s Major Splice Donor (MSD) site (HXB2 coordinates 743-745), suggesting it could be replication impaired (**Figure 2B**). Since most documented MSD deletions are larger (>20 bases) (3, 22), and encompass both the MSD and the cryptic “GT” splice donor site 4 bases downstream (20) (which is preserved in the NSV provirus), we investigated the impact of this small deletion in the context of the participant’s unique recombinant virus. As our near-full-length proviral sequences did not capture HIV’s long terminal repeats (LTRs), we single-genome-amplified proviral sequences spanning the 5’ LTR into *gag* (**Figure 3A, 3B**), and selected those that best matched our proviruses of interest to construct three full-length autologous molecular clones: the NSV genome harboring the three-base MSD deletion (“NSV”), a modified NSV genome with the MSD deletion repaired (“NSV rescue”) and a representative intact provirus from the participant (“C14 control”) (**Figures 3C, 3D**). Transfection of HEK-293T cells with equal plasmid copy numbers revealed that the NSV molecular clone produced virions, though to a three-fold lesser extent compared to NSV rescue (p=0.006) and C14 control (p=0.01), while virion production from the latter two clones was comparable (p=0.2) (**Figure 4A**). This indicated that the three-base MSD deletion reduced, but did not prevent, virion production, which was consistent with the observation of NSV *in vivo*.

**Figure 3.**
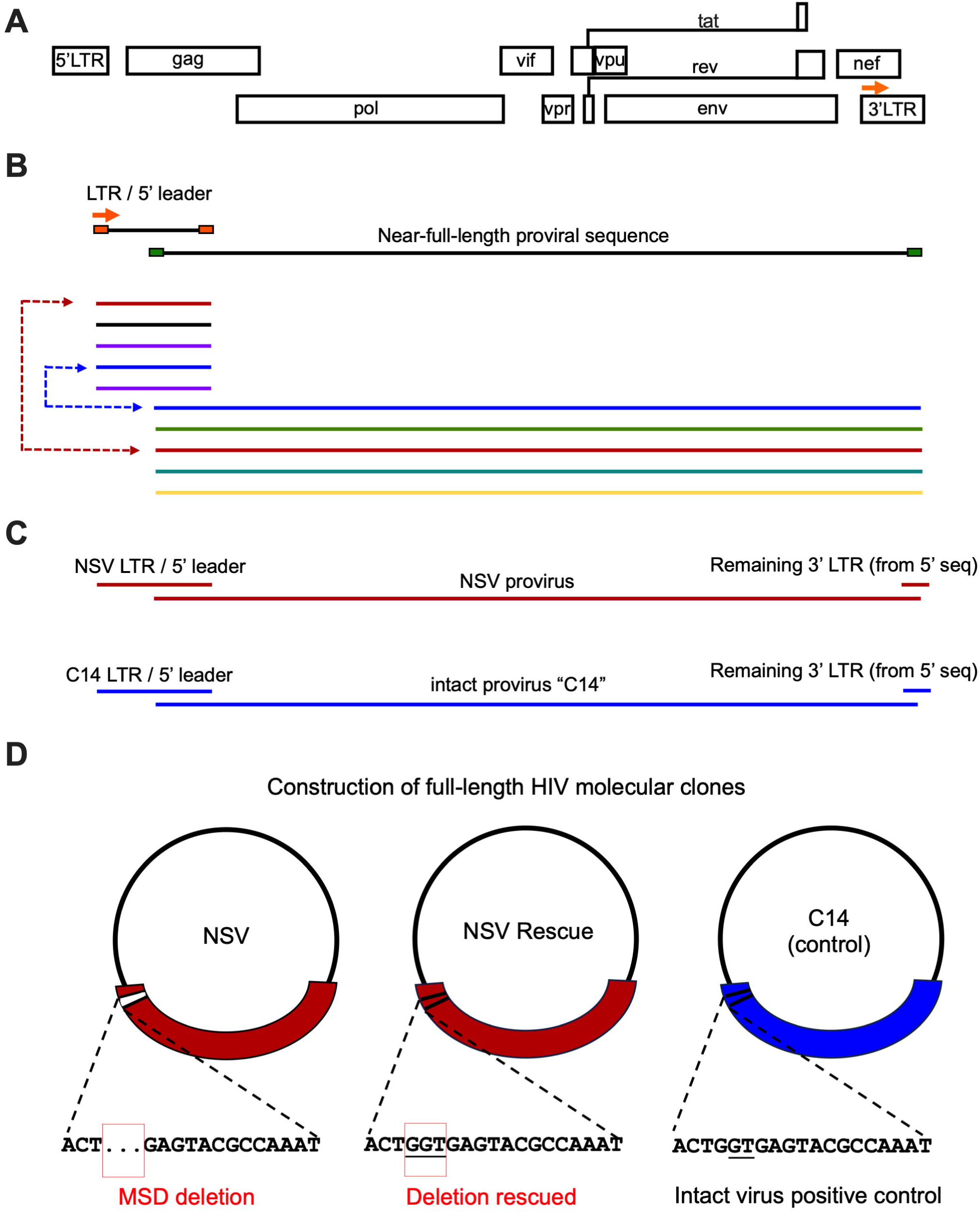
PCR amplification and construction of three autologous molecular clones. (A, B) Using autologous primers that bound to the beginning of the participants’ LTR (designed based on the 3’ LTR; orange arrow) and *gag* respectively, we isolated 81 LTR/*gag* sequences by single-genome amplification. (C, D) The LTR sequences that best matched each near-full-length provirus of interest was used to construct three full-length molecular clones: the NSV provirus (“NSV”), the NSV provirus with the three-base deletion rescued (“NSV rescue”) and another intact provirus from the participant as a control (“C14 control”). The MSD region of each molecular clone is shown below each plasmid.

**Figure 4.**
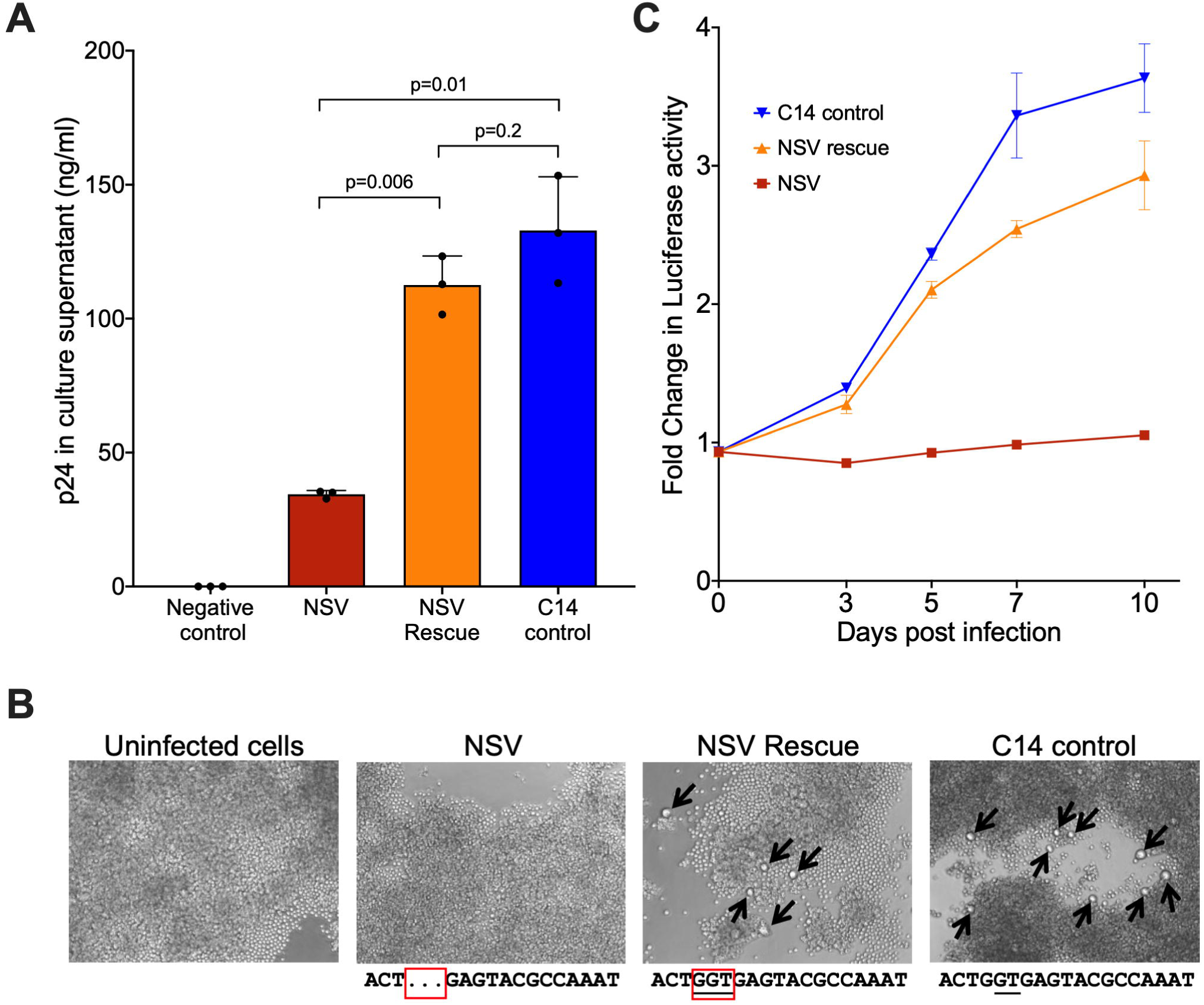
Assessing virion production and replication capacity of the HIV molecular clones. (A) The ability of each molecular clone to produce virions was assessed by transfecting equal amounts of plasmid into HEK-293T cells and quantifying p24 in culture supernatants 48 hours later. Bars and whiskers represent the mean and standard deviation of three technical replicates. *P*-values were calculated using an unpaired t-test with Welch’s correction. (B) Representative microscopy images showing the formation of syncytia (black arrows) in the NSV-rescue and C14-infected, but not NSV-infected cultures, at 10 days post-infection. The MSD region sequence of each virus is shown below each image. (C) Viral spread in culture over 10 days, assessed by quantifying Gaussia luciferase in culture supernatants. Data points and whiskers represent the mean and standard deviation of three technical replicates, respectively.

We next investigated the NSV clone’s ability to replicate by infecting Sup-GGR reporter cells with equal amounts of NSV, NSV rescue and C14 virus (inoculum normalized by p24^Gag^ levels). Syncytia were observed in the NSV rescue and C14 cultures after 3 days, while the NSV culture resembled uninfected cells over 10 days (**Figure 4B**). Consistent with this, the NSV rescue and C14 cultures showed evidence of viral spread as measured by luciferase production, whereas the NSV culture did not (**Figure 4C**). This indicates that the MSD deletion conferred a profound replication impairment that could be restored upon deletion repair.

### Identifying the cell type harboring the NSV provirus

Because the three-base MSD deletion was essentially exclusive to the NSV provirus clone (only two other proviruses encoded it, representing 0.5% of the overall proviral pool), we exploited it as a molecular feature to distinguish the NSV from other proviruses. We developed a duplex droplet digital PCR (ddPCR) assay that targeted HIV’s MSD and envelope regions, where the MSD-region probe matched the NSV provirus but was still able to bind, albeit less well, to other autologous proviruses (**Figure S3A**). Using synthetic DNA templates, we demonstrated that the MSD probe produced a high-amplitude signal for the NSV sequence and a markedly lower-amplitude signal for other autologous sequences, allowing clear discrimination of the NSV sequence even in mixed templates (**Figure S3B**). We further confirmed that the duplexed assay clearly distinguished the NSV provirus from others using our three molecular clones (**Figure S3C**). Application of this assay to DNA extracted from 792,000 of the participant’s blood CD4+ T cells revealed that the NSV provirus was present at 32 copies/million CD4+ T cells (**Figure 5A and S4A**).

**Figure 5.**
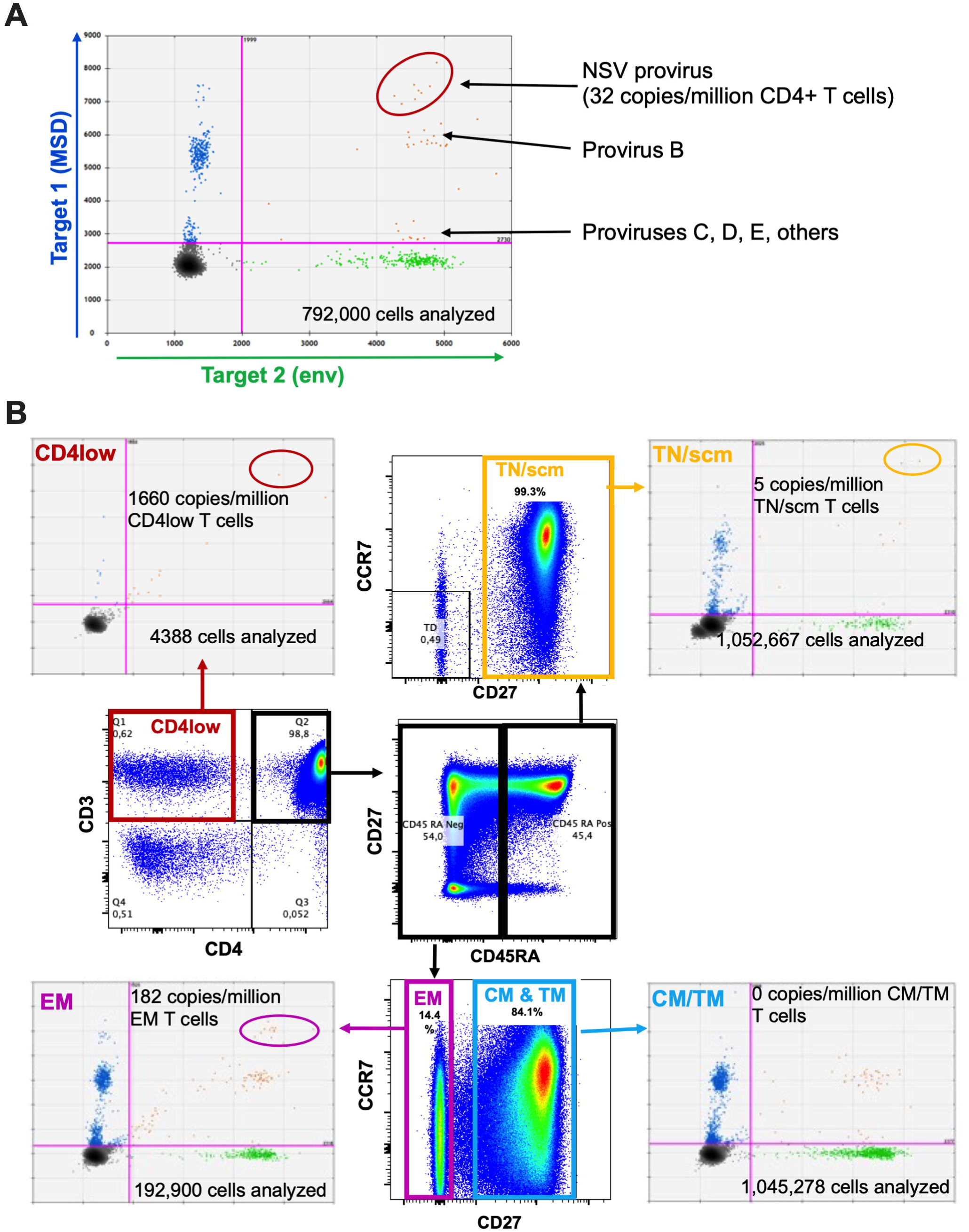
The NSV provirus is found in multiple CD4+ T-cell subsets but is most abundant in effector memory-like CD4+ T-cells. (A) Representative ddPCR data from the analysis of bulk genomic DNA from 792,000 CD4+ T-cells revealed that the NSV provirus was present at an estimated 32 copies per million total CD4+ T-cells. (B) The flow cytometry plots in the interior of the figure depict the sorting strategy for blood CD4+ T-cell subsets. Blood CD4+ T-cells were isolated by negative selection and then sorted into: Naïve or stem cell memory cells (TN/scm; CD45RA+CD27+; boxed in yellow in the top flow plot); Central or Transitional Memory cells (CM/TM; CD45RA-CD27+; boxed in blue in the bottom flow plot); Effector Memory-like cells (EM; CD45RA-CD27-; boxed in purple in the bottom flow plot); and cells expressing low levels of CD4 (CD4low; boxed in red in the leftmost flow plot). Matching colored arrows point to representative raw ddPCR data from each subset, with estimated NSV provirus copy numbers and total number of cells analyzed shown for each subset.

We next applied this assay to DNA extracted from different blood CD4+ T-cell subsets to identify those that harbored the NSV provirus. In our initial attempt, we sorted naïve/stem cell memory (TN/SCM), central memory (CM), transitional memory (TM), and effector memory (EM) subsets based on CD45RA, CD27, and CCR7 expression (**Figure S4B**). Despite extracting DNA from >1 million TN/SCM cells and ∼500,000 TM cells, only one double-positive event representing the NSV provirus was detected in each of these subsets, and no NSV-specific events were detected in CM and EM cells. This indicated that this sorting strategy had failed to capture the NSV provirus-harboring cell subset.

We repeated using a more inclusive sorting strategy, capturing TN/SCM-like cells (defined as CD45RA+/CD27+), TCM/TM (CD45RA-/CD27+), and TEM-like (CD45RA-/CD27-) subsets (**Figure 5B**), where the TEM-like gate now included the participant’s unusual CD45RA-CD27-CCR7+ cell population that was previously excluded (see **Figure S4B**). Reasoning that the NSV provirus might be expressing functional HIV accessory proteins, including Nef that can downregulate CD4, we also sorted T-cells that displayed reduced surface CD4 expression (“CD4low” subset). Using this strategy, we abundantly detected the NSV provirus in the TEM-like subset, at a frequency of 182 copies/million cells, suggesting the participant’s unusual CD45RA-CD27-CCR7+ cell population is the subset that predominantly harbors this provirus. Consistent with our initial sort, we also detected two events representing the NSV provirus in the TN/SCM-like subset, though none in the TCM/TM-like subset, despite analyzing >1 million cells for each of these subset. We also detected one NSV provirus event in CD4low cells, despite analyzing fewer than 4400 cells in this subset, indicating that cells harboring the NSV provirus can downregulate CD4.

### The NSV provirus exhibits an immune-evasive HIV protein expression profile

We hypothesized that the reason the NSV provirus clone had produced viremia for so long without being eliminated was because it exhibited an HIV protein expression profile that allowed it to evade immune detection. Based on evidence that HIV subtype B proviruses with 5’ defects produced non-infectious virions with decreased Env incorporation (3), we investigated cell-surface Env in NSV-provirus-expressing cells, reasoning that such defects could explain both impaired virion replication and clone persistence due to reduced *env* expression protecting NSV provirus-expressing cells from elimination by antibody-dependent cellular cytotoxicity (ADCC) (27–33). After transfecting HEK-293T cells with equal copy numbers of the three molecular clones, we observed that NSV-transfected cells that were p24+ expressed substantially less surface Env compared to NSV rescue and C14-transfected cells (∼10% of p24+ NSV-transfected cells were Env+, compared to >40% for NSV rescue and C14) (**Figure 6**). This confirmed that Env expression is impaired in the NSV provirus, and this defect is rescued by restoring the MSD deletion.

**Figure 6.**
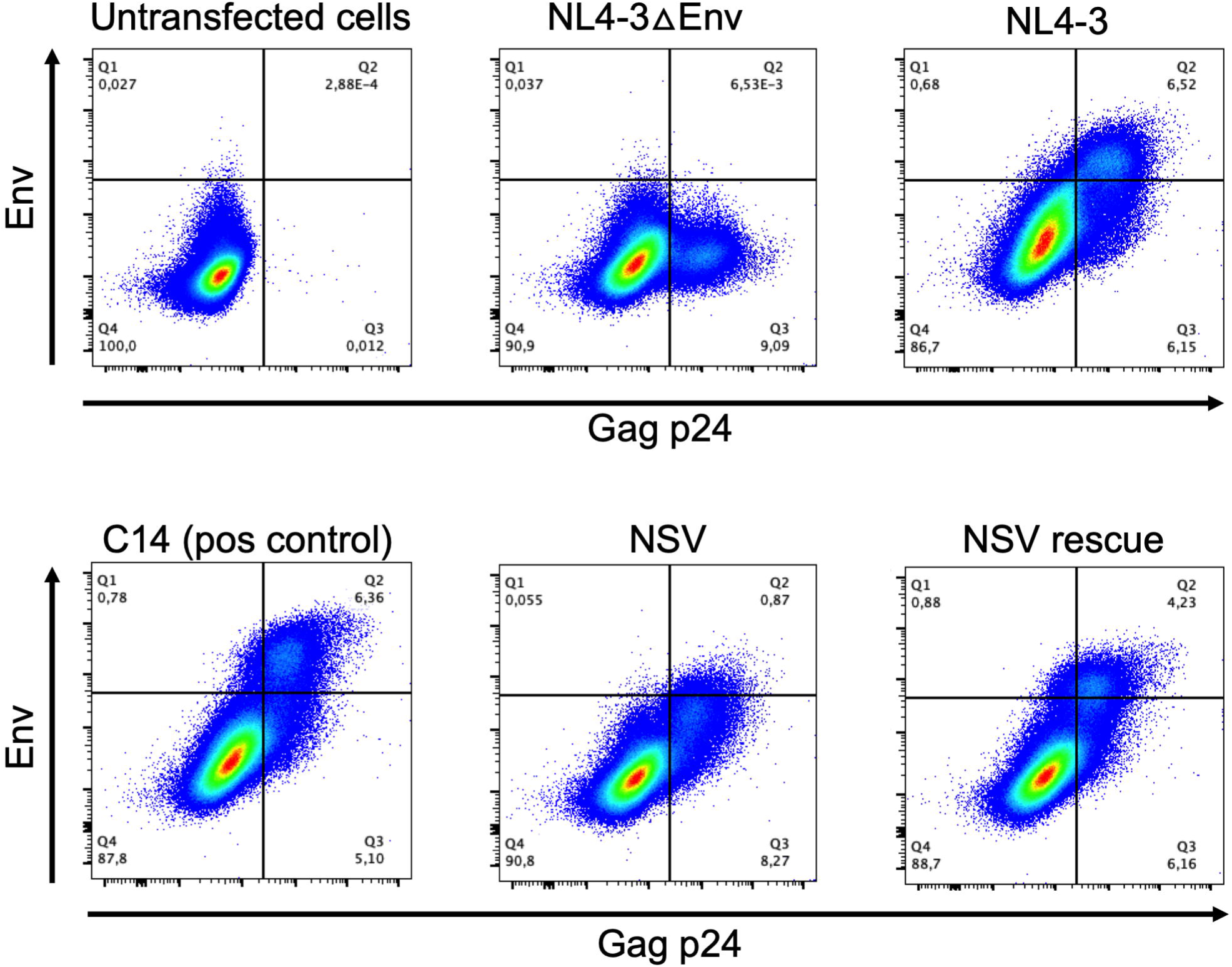
The NSV virus displays impaired envelope expression. Representative flow cytometry plots showing intracellular p24 and cell-surface Env expression in HEK-293T cells 42 hours post-transfection with control HIV molecular clone plasmids (top row), and participant molecular clones (bottom panel). Data from one of two independent experiments is shown.

We also investigated the NSV virus’s ability to downregulate HLA class I, which would allow infected cells to evade detection by CD8+ cytotoxic T cells. We transfected HEK-293T cells with equal copy numbers of the NSV, NSV rescue and C14 molecular clones, and assessed HLA-A*02 cell surface expression by flow cytometry (**Figure 7A**). All three viruses exhibited similar HLA downregulation abilities, strongly suggesting that, despite its MSD deletion, the NSV virus produces functional Nef. Using the same transfection approach, we confirmed comparable Nef expression by the three molecular clones by flow cytometry (**Figure 7B**).

**Figure 7.**
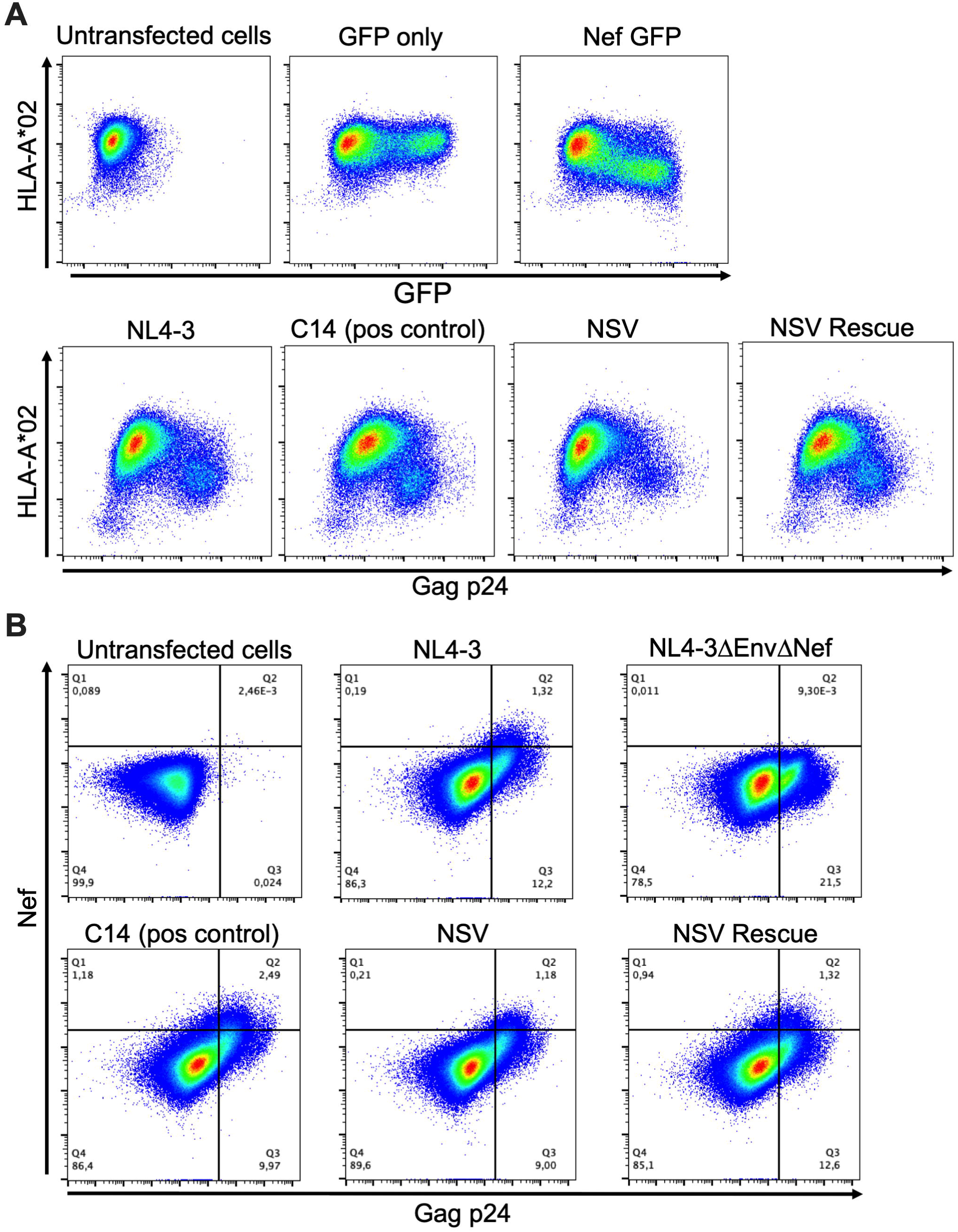
The NSV robustly downregulates HLA class I and expresses Nef. (A) Top row: Representative flow cytometry plots showing cell-surface HLA-A*02 expression and intracellular GFP expression in HEK-293T cells transfected with control plasmids, including empty pSELECT-GFP (negative control) and pSELECT-GFP expressing *nef* from the HIV subtype B reference strain SF2 (positive control). Bottom row: Representative flow cytometry plots showing cell-surface HLA-A*02 and intracellular p24 expression in HEK-293T cells transfected with a full-length NL4-3 molecular clone (positive control) or participant molecular clones. Data from one of three independent experiments is shown. (B) Representative flow cytometry plots showing intracellular Nef expression in HEK-293T cells transfected with control molecular clones (top row) or participant molecular clones (bottom row).

### The NSV virus uses two alternative D1 splice donor sites

Given that the NSV virus expressed Nef, and to a lesser extent Env, we investigated what alternative splice donor (D1) site was being used to produce these, and potentially other, spliced HIV transcripts. Using primers specific for the NSV sequence (**Table S1**), we single-genome amplified 61 spliced transcripts from HEK-293T cells transfected with the NSV molecular clone. Of these, 57 (93%) used a novel cryptic GT motif located 31 bases upstream of the MSD, which we called D1*, while four (7%) used a previously described cryptic GT motif located four bases downstream of the MSD, which we called D1** (3, 20) (**Figure 8A**). Of note, D1* fell within HIV’s palindromic Dimerization Initiation Site (DIS) sequence (34, 35), which was G<u>GT</u>ACC in this participant’s HIV population (D1* underlined). This DIS sequence differs markedly from the canonical GCGCGC (in subtypes B and D) or GTGCAC (in other subtypes) (36, 37).

**Figure 8.**
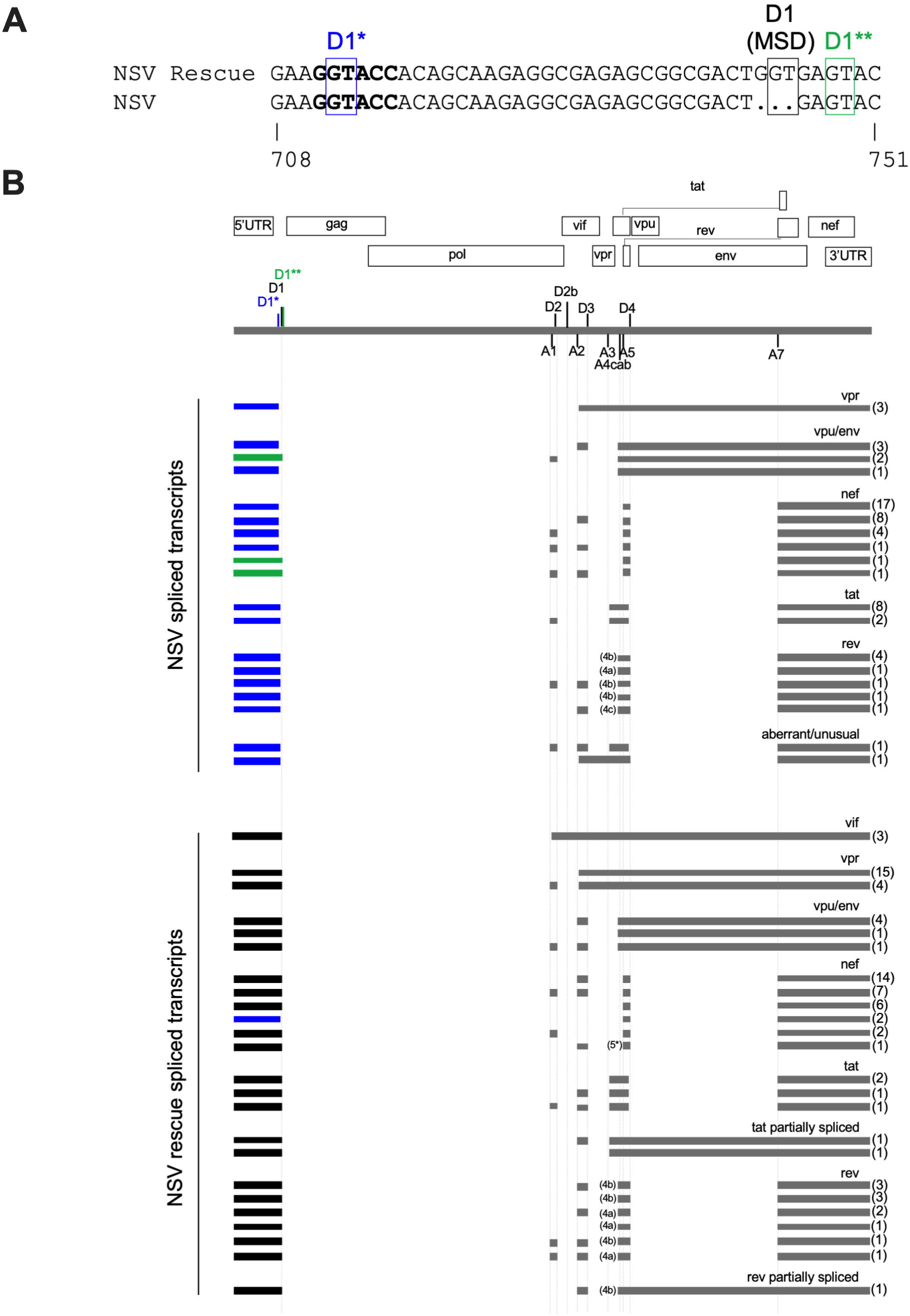
The NSV provirus uses two alternative RNA splice donor sites, but overall spliced transcript composition is nevertheless altered. (A) Nucleotide alignment of NSV and NSV rescue proviruses, showing the MSD (black) and the two alternative splice donor sites used by the NSV provirus: D1* (31 bases upstream; blue) and D1** (4 bases downstream; green). HIV’s Dimerization Initiation Site (DIS) is shown in bold. HXB2 genomic coordinates are shown below the alignment. (B) Genomic map of the NSV provirus, showing the locations of major splice donor and acceptor sites. Individual spliced HIV transcripts detected in cells transfected with equal amounts of NSV and NSV rescue molecular clones are shown below, where the numbers in parentheses denote how many times each transcript was observed. Exons are depicted as horizontal bars, with the first exon colored based on the D1 site used: MSD (black), D1* (blue) or D1** (green). For the splice acceptor sites 4c, a and b, the specific site used is shown to the left of this exon. One *nef* transcript observed in NSV rescue-transfected cells used an alternative A5 acceptor site, labeled 5*.

In contrast, yet as expected, 76 of 78 (97%) of spliced transcripts from cells transfected with the NSV rescue clone used the MSD site, though two transcripts (3%) encoding Nef used the cryptic D1* site within the DIS. Though NSV-transfected cells produced a variety of spliced transcripts from the 1.8kb and 4kb classes, their composition was subtly different from those of NSV rescue-transfected cells (**Figure 8B**). First, *vpr*-encoding transcripts were significantly less abundant in NSV-transfected compared to NSV rescue-transfected cells (p=0.0019). Furthermore, *vif*-encoding transcripts were not detected in NSV-transfected cells, though *vif* is typically the least-abundant spliced HIV transcript (38, 39). Unusual/aberrant transcripts were also detected in NSV-transfected cells, but not in NSV rescue-transfected cells. Multiply spliced transcripts encoding *tat*, *rev* and *nef* from NSV-transfected cells also had one fewer exon on average: a median of three, compared to a median of four in cells transfected with the NSV rescue plasmid (p=0.0015).

Most strikingly, for all classes of spliced transcripts, RNA extracts from NSV rescue-transfected extracts had to be diluted ∼10-fold more than NSV-transfected extracts to achieve limiting dilution, even though cultures were transfected with the same amount of plasmid, and RNA was extracted from the same number of cells. This strongly suggested that use of alternative D1 sites markedly reduced spliced transcript production. Using RT-ddPCR, we confirmed that 1.8kb-class HIV transcripts were 13 times less abundant in NSV-compared to NSV-rescue-transfected cells (p=0.0006) (**Figure S6**). Specifically, in NSV-rescue transfected cells, 1.8kb-class transcripts outnumbered housekeeping transcripts by a median of 382 (Interquartile range [IQR] 361-397)-fold, compared to only median of 29 (IQR 23-32)-fold in NSV-transfected cells (**Figure S6**). Together, these observations indicate that the use of alternative D1 sites markedly reduced spliced transcript abundance and altered their composition, helping to explain some of the defects observed.

### The NSV spontaneously resolved after contraction of the clone population

The participant’s NSV spontaneously resolved after 4.5 years. Analysis of 1.7 million CD4+ T cells shortly after NSV resolution by ddPCR revealed that the NSV provirus population had diminished from the estimated 32 copies/million CD4+ T-cells initially measured during the viremic period, to an estimated four copies/million CD4+ T cells after viremia resolution, an 8-fold reduction (**Figure S5**). This is consistent with the NSV resolving upon contraction of the NSV-provirus harboring clonal population.

## DISCUSSION

We describe a case of non-suppressible viremia (NSV) that lasted ∼4.5 years in an individual living with a non-B HIV subtype. The viremia was caused by a clonally expanded provirus harboring a three-base deletion in HIV’s major splice donor (MSD) site, which impaired virus replication. The virus compensated for the MSD deletion by using two alternative D1 splice donor sites: one located four bases downstream, previously described in subtype B (3, 20) and another novel site located 31 bases upstream within this individual’s distinctive HIV dimerization initiation site (DIS) sequence. These observations highlight the plasticity of splice donor site usage in HIV (21, 40, 41) and underscore the importance of including diverse viral isolates in HIV persistence research.

Despite using alternative D1 sites, the three-base MSD deletion nevertheless markedly impacted spliced HIV transcript abundance and composition, providing insights into the virus’s replication impairment. Similar to a recent NSV case in HIV subtype B that was also caused by an MSD-deleted provirus (3), the replication impairment in the present case was attributable, at least in part, to impaired Env expression. The underlying mechanism of reduced Env expression, however, was different. In the published case, which featured a >20 base MSD deletion, Env expression was defective because spliced transcripts encoding *rev* were not produced (*env* transcripts require Rev for nuclear export) (3, 29, 42). In contrast, the present NSV provirus produced *rev* transcripts. Instead, we speculate that poor Env expression on NSV-transfected cells resulted from a >10-fold reduction in the overall abundance of spliced transcripts caused by the MSD deletion. We also noted that the bicistronic *vpu*/*env*-encoding transcripts in NSV-transfected and NSV rescue-transfected cells differed subtly in their exonic composition, which might influence which of the two genes is expressed: for example, half of *vpu*/*env*-encoding transcripts in NSV-transfected cells lacked the upstream exon between the A2/D3 splice sites, whereas all but one *vpu*/*env*-encoding transcript in NSV rescue-transfected cells retained this exon.

Importantly, cells expressing the NSV provirus exhibited an HIV protein expression profile that almost certainly helped these cells evade immune detection despite prolonged viremia production *in vivo*. Though NSV-provirus-expressing cells exhibited markedly reduced spliced transcript levels, they still expressed sufficient functional Nef protein to downregulate cell-surface HLA class I, which would have enabled them to evade CD8+ cytotoxic T-lymphocyte (CTL)-mediated elimination despite HIV protein production (43, 44). At the same time, reduced cell-surface Env levels would protect NSV-provirus-expressing cells from elimination by antibody-dependent cellular cytotoxicity (ADCC). In fact, downregulation of CD4 (*e.g.* by Nef or Vpu) would have further enhanced the resistance of NSV provirus-expressing cells to ADCC by preventing the required Env-CD4 interaction on the cell surface (33, 45–48). Our detection of the NSV provirus *ex vivo* in a CD4+ T-cell that had downregulated its CD4 receptor also supports this notion. On the other hand, the fact that most NSV-provirus-harboring cells expressed normal CD4 levels (as demonstrated by the cell sorting data, which detected CD4 on all NSV-provirus-expressing cells except the aforementioned one) suggested that only a fraction of NSV-harboring clones were actively producing HIV transcripts/proteins at any given time. Viremia production by only a fraction of the expanded clone population is also consistent with previous reports (3, 18).

Though HIV can persist in all CD4+ T-cell subsets, including naïve and stem cell memory cells (49–53), proviruses are generally enriched in more differentiated memory subsets, with expanded clones most likely to be effector memory cells (54–56). We detected the NSV provirus in three CD4+ T-cell subsets: at low levels in naive/stem cell memory (SCM) and transitional memory cells (5 and 7 copies per million cells, respectively) and at high levels in effector memory (EM) T-cells (182 copies/million cells). We posit that the NSV virus’s long-term persistence *in vivo* is likely due to it’s immediate ancestor infecting a T_SCM_, during which the three-base MSD deletion was introduced during reverse transcription (we infer that this initial infection occurred in T_SCM_ because the NSV virus is predicted to use CCR5, which is expressed by T_SCM_ but not naïve T cells). This cell then differentiated into other memory cell types, including an atypical EM subset that expressed a CD45RA-CD27-CCR7+ phenotype, which was the likely source of prolonged viremia. After 4.5 years, the NSV-harboring clonal population contracted ∼8-fold in size, leading to the spontaneous resolution of the NSV. This is consistent with the “waxing and waning” of proviral clones on years-long timescales (57).

This study has some limitations. The events that triggered the NSV and its resolution are unknown. A recent paper suggested that changes in HIV therapy − particularly a switch from a non-nucleoside reverse transcriptase inhibitor to an integrase inhibitor − could temporarily perturb the reservoir in some individuals (58). Though it is unclear whether therapy changes played a role in the present case, the onset of NSV roughly coincided with a switch to a bictegravir-containing regimen (from an elvitegravir-containing one), and resolved approximately one year after bictegravir was removed from the regimen. Due to insufficient cell numbers, we could not further phenotypically characterize the atypical CD45RA-CD27-CCR7+ CD4+ T cell subset that likely produced the viremia, nor determine the antigen specificity of this clone, though a recent report suggested that CD4+ T-cells that are reactive to self-antigens can contribute to NSV (59). Potentially consistent with this, the participant experienced temporary cognitive impairment coincident with NSV onset that did not appear to be explained by virus presence in cerebrospinal fluid (no HIV was detected in CSF collected during this time). Since we did not determine the integration site of the NSV provirus, we cannot definitively state that it is a clone, though the likelihood is extremely high, as identical proviruses shared integration sites in 100% of cases in a recent study (60). Finally, although the NSV virus was profoundly replication impaired *in vitro*, we cannot definitively say it is fully replication incompetent. In fact, we isolated two proviruses that were near-identical to the NSV provirus (one differed by only a single base; the other by two). Given the number of proviruses sequenced, these mutations are broadly within our assay error estimate of 1.8×10^-5^ mutations per base sequenced (61), but we cannot rule out that they arose through replication of the MSD-deleted virus *in vivo*.

HIV cure efforts focus on eliminating reservoir cells, specifically defined as those harboring genetically intact proviruses capable of producing infectious virus. This category excludes proviruses with MSD mutations or deletions, which are assumed to be defective (62–65). The possibility that some proviruses with MSD deletions might be able to replicate (albeit poorly) *in vivo* has potential implications for HIV cure strategies as well as transmission risk during prolonged NSV, and underscores the need to understand this class of proviruses better. But even if the *in vivo* replication competence of MSD-defective proviruses is effectively zero, our results add to the growing evidence that they can clonally expand (66), produce viral transcripts (23), HIV proteins (41, 67), and virions over long periods (3), and can exhibit HIV protein expression profiles that facilitate immune evasion and persistence. These functions suggest that these proviruses are likely to be biologically and clinically relevant. Given that HIV proteins produced by defective proviruses likely drive inflammation (68, 69) and have been associated with clinical outcomes including nonresponse to ART (70), and given the recent demonstration that defective proviruses can conditionally replicate upon superinfection with intact HIV (71), a better understanding of this abundant yet underappreciated proviral class is needed. And, given the unique genetic features of the present case (e.g. 3-base MSD deletion; novel splice donor site within a distinctive DIS), this research should include diverse, globally representative HIV isolates.

In conclusion, our findings confirm that NSV originates from clonally expanded proviruses with MSD defects (3, 4, 17). Currently, among the major HIV treatment guidelines, only the European ones acknowledge NSV due to cellular proliferation, but only in the specific context of persistent viremia between 50-200 copies/ml in persons fully adherent to ART and in the absence of drug resistance (24). All guidelines continue to recommend that plasma viral loads >200 HIV RNA copies/mL during ART be managed as virologic failure with regimen modification or intensification (6, 7), even though NSV, which can exceed 200 copies/mL as demonstrated by us and others (3, 4, 17), is not resolvable by this action (and in fact therapy modification would expose individuals to unnecessary drug toxicities, not to mention anxiety when the viremia fails to resolve). Clinical recommendations should be developed to help discriminate persistent viremia that is due to drug adherence, pharmacokinetic issues or drug resistance development (which is clinically actionable), from NSV (which is not). Presumptive NSV cases could be identified by ruling out suboptimal adherence, pharmacological interactions and new drug resistance, where the suggested management would be to maintain the current ART regimen, continue to monitor viral load, CD4 T-cell counts and co-morbidities, repeat drug resistance testing at intervals, and if possible undertake more in-depth HIV RNA sequencing to detect clonality and MSD defects, where the latter would also help assess potential transmission risk during prolonged NSV (72). Indeed, the findings of the present study reassured both participant and care provider that the NSV was not caused by any action (or lack thereof) on their part, allowing the provider to de-intensify ART and alleviate concerns regarding transmission risk during this period.

## METHODS

### Ethics statement

This study was approved by the Providence Health Care/University of British Columbia and Simon Fraser University research ethics boards. The participant provided written informed consent.

### Amplification and sequencing of plasma HIV RNA

Our clinically accredited laboratory at the BC Centre for Excellence in HIV/AIDS performs HIV drug resistance genotyping for nearly all of Canada, where the genotyped region covers HIV protease and codons 1-400 of reverse transcriptase (73). Briefly, total nucleic acids were extracted from 500 μL of plasma on a NucliSENS EasyMag (bioMerieux) after which bulk cDNA was generated using an HIV-specific reverse primer and NxtScript Reverse Transcriptase (Roche). Subsequent nested PCR reactions were performed using the Expand™ High Fidelity PCR System (Roche). Partial 5’ leader/gag and *gp41* regions were also isolated from plasma HIV RNA by single-genome amplification. For this, nucleic acid extracts were DNAse-treated prior to amplification and cDNA was endpoint-diluted such that subsequent nested PCR reactions yielded <30% positive amplicons. All PCR primers are listed in **Table S1**. Amplicons were sequenced using a 3730xl Automated DNA Sequencer (Applied Biosystems) and chromatograms were analyzed using Sequencher v.5.0 (Gene Codes). HIV drug resistance interpretations were performed using the Stanford HIV drug resistance database (74).

### Near-full-length HIV proviral sequencing

Peripheral blood mononuclear cells (PBMC) were isolated from whole blood by standard density gradient separation and cryopreserved at –150°C until use. CD4+ T-cells were isolated from PBMC by negative selection (STEMCELL Technologies) and genomic DNA was extracted using the QIAamp DNA Mini kit (Qiagen). Near full length, single-genome HIV proviral sequencing was performed as previously described (61). Briefly, genomic DNA was diluted such that <30% of resulting nested PCR reactions, performed using Platinum Taq DNA Polymerase High Fidelity (Invitrogen), yielded an amplicon (**Table S1**). Amplicons were sequenced on an Illumina MiSeq and reads were *de novo* assembled using an in-house modification of the Iterative Virus Assembler (75) implemented in the custom software MiCall (version 7.17.0) (http://github.com/cfe-lab/MiCall) to generate a consensus sequence. The genomic integrity of sequenced proviruses was determined using an in-house modification of the open-source software HIV SeqinR (63), where an intact classification required all HIV reading frames, including accessory proteins, to be intact. Sequences with 100% identity across the entire amplicon were considered identical and clonal. HIV subtyping was performed using the recombinant identification program (RIP) hosted on the Los Alamos HIV sequence database web server (76) using a window size of 400 and a confidence interval of 90%.

### Phylogenetic analysis

HIV sequences were codon-aligned using HIVAlign (MAFFT option) (77) hosted on the Los Alamos HIV sequence database (78) and manually edited using AliView (79). Maximum likelihood phylogenies were constructed using IQ-TREE 2 (80) following automated model selection with ModelFinder (81) using an Akaike information criterion (AIC) (82). Phylogenies were visualized using the R package ggtree (83).

### HIV molecular clone construction and replication capacity assessment

We constructed full-length HIV molecular clones in a pUC57Brick vector (GenScript), representing the NSV provirus (“NSV”), the NSV provirus with the three-base MSD deletion rescued (“NSV rescue”) and another representative intact provirus from the participant (“C14”), where the autologous HIV LTR and 5’ leader regions were reconstructed by single-genome amplifying sequences from proviral genomic DNA (primers in **Table S1**) and selecting the one that best matched each near-full-length provirus in the overlap region (**Figure 3**). Molecular clones were propagated in recombinase-deleted Stbl3 *E. coli* (Invitrogen). To generate virus stocks, 13 µg of plasmid was transfected into 2.5 million HEK-293T cells using lipofectamine LTX (Thermo Fisher Scientific). Supernatants were collected 48 hours later, and p24 levels were measured by ELISA (XpressBio). Virus replication was assessed using the Sup-GGR reporter cell line (84), which contains a Tat/Rev-dependent expression cassette that produces humanized *Renilla* GFP and *Gaussia* luciferase upon HIV infection. For each virus type, we infected one million cells with 13.8 ng of p24. Syncytia formation was monitored by light microscopy, and supernatant *Gaussia* luciferase levels were quantified using a bioluminescence assay (Pierce^TM^ Gaussia Luciferase Glow, Thermo Scientific) for 10 days.

### Quantification of the NSV provirus using ddPCR

We designed a dual-target droplet digital PCR (ddPCR) assay modeled upon the Intact Proviral DNA assay (IPDA) (85, 86) to discriminate the NSV provirus from the participant’s other proviruses based on its unique MSD deletion. The assay’s 5’ (MSD) target combined the published IPDA primers with a custom probe that matched the NSV provirus sequence in the MSD region (**Table S1** and **Figure S3**). The assay’s 3’ (*env*) target used a published secondary *env* probe (86) with autologous primers (**Table S1**). Reactions were prepared as previously described (85, 86) and data were collected on a QX200 Droplet Reader (BioRad), analyzed using QuantaSoft software (BioRad, version 1.7.4). The 5’ (MSD) target was validated using synthetic DNA templates (IDT gBlocks) spanning HXB2 coordinates 669-919, representing the NSV provirus and the four next most abundant proviruses (**Figure S3**). At least four technical replicates were performed for each ddPCR reaction and merged for analysis.

### Sorting of CD4+ T-cell subsets

We isolated CD4+ T cell subsets from a minimum of 400 million cryopreserved PBMC as previously described (87) with some modifications. Total CD4+ T cells were labeled with 7-Aminoactinomycin D (AAD) live/dead dye, Peridinin-Chlorophyll-Protein (PerCP)-labeled mouse anti-human CD3 antibody, fluorescein isothiocyanate (FITC)-labeled mouse anti-human CD45RA antibody, and Brilliant Violet (BV421)-labeled mouse anti-human CD27 antibody (all from Biolegend), and allophycocyanin (APC)-labeled mouse anti-human CD4 antibody and phycoerythrin (PE)-Cy7-labeled rat anti-human CCR7 antibody (both from BD Biosciences). Naive/stem cell memory (T_N/SCM_), central memory (T_CM_), transitional memory (T_TM_), and effector memory (T_EM_) CD4^+^ T cells were separated using a FACSAria cell sorter (BD Biosciences) according to the strategy shown in **Figures 5** and **S4**.

### Flow cytometric analysis of HIV protein expression and HLA downregulation

To measure cell-surface HIV Env expression, we transfected 2.5 million HEK-293T cells with 13µg of NSV, NSV rescue, C14, or control plasmid (NL4-3 or NL4-3Δenv). Forty-two hours later, cells were surface-stained using a cocktail of three primary Env antibodies, 3BNC117 (NIH HIV Reagent program; ARP-12474), VRC01 (ARP-12033), and PGT128 (ARP-13352) (88–91), and PE-labeled anti-human IgG (Biolegend) as secondary. Cells were then stained with Zombie Aqua live/dead dye (Biolegend), fixed/permeabilized (cytofix/cytoperm reagent; BD Biosciences) and intracellularly stained with an APC-labeled anti-p24 antibody (28B7 clone, MediMabs, MM-0289-APC). To measure HLA-A*02 downregulation, we transfected 2.5 million HEK-293T cells with 13µg of NSV, NSV rescue or C14 plasmid. A pSELECT GFP-reporter plasmid expressing HIV Nef and an empty pSELECT vector were used as controls. Twenty-six hours later, cells were stained with Zombie Aqua live/dead dye (Biolegend) and APC-labeled anti HLA-A*02 antibody (BB7.2 clone, Biolegend). After fixation/permeabilization, cells were intracellularly stained with PE-labeled anti-p24 antibody (KC57 clone, Beckman Coulter). To measure intracellular HIV Nef expression, we transfected 2.5 million HEK-293T cells with 13µg of NSV, NSV rescue, C14, or control plasmid (NL4-3 or NL4-3ΔEnvΔNef). Forty-two hours later, cells were stained with Zombie NIR live/dead dye (Biolegend), fixed/permeabilized, and intracellularly stained for Nef using rabbit polyclonal anti-HIV-1 Nef serum as primary (NIH HIV Reagent program, ARP-2949) and BV421-labeled donkey anti-rabbit IgG as secondary (Biolegend). Gag p24 was detected as described above using the APC-labeled 28B8 clone (MediMabs). Flow cytometry was performed on a Cytoflex S instrument (Beckman Coulter).

### Characterization and quantification of spliced HIV transcripts

We transfected HEK-293T cells with 13µg of NSV or NSV rescue plasmid and isolated total cellular RNA 24 hours later (PuroSPIN™ Total RNA Purification Kit; Luna Nanotech). We generated cDNA using reverse primers designed to capture 4kb-class, 1.8kb-class and universal spliced HIV transcripts, and amplified these using single-genome approaches (primers in **Table S1**). Amplicons were Sanger-sequenced as described above and analyzed in Sequencher v.5.0 (Gene Codes). To quantify the abundance of 1.8kb-class HIV transcripts, we designed a duplex RT-ddPCR assay that simultaneously measured these, along with a human housekeeping transcript (RPP30). Specificity for the 1.8kb HIV transcript class was ensured by placing the probe across the D4/A7 splice junction (primers in **Table S1**).

### Statistical analyses

Statistical analyses were performed in Prism (version 8.4.3).

## Data availability

GenBank submission is in progress.

## Acknowledgements

We thank Maria Velasquez, Bruce Ganase and Landon Young for assistance with participant recruitment and phlebotomy, and Don Kirkby for assistance with proviral genome assemblies. We are grateful to Drs. Nicolas Chomont and Rémi Fromentin for their helpful guidance on sorting CD4+ T cell subsets. We thank the BC Centre for Excellence in HIV/AIDS clinical laboratory team for support. We gratefully thank the study participant and their HIV care provider, without whom this research would not have been possible.

## Legends for Supplementary Figures

**Figure S1.**
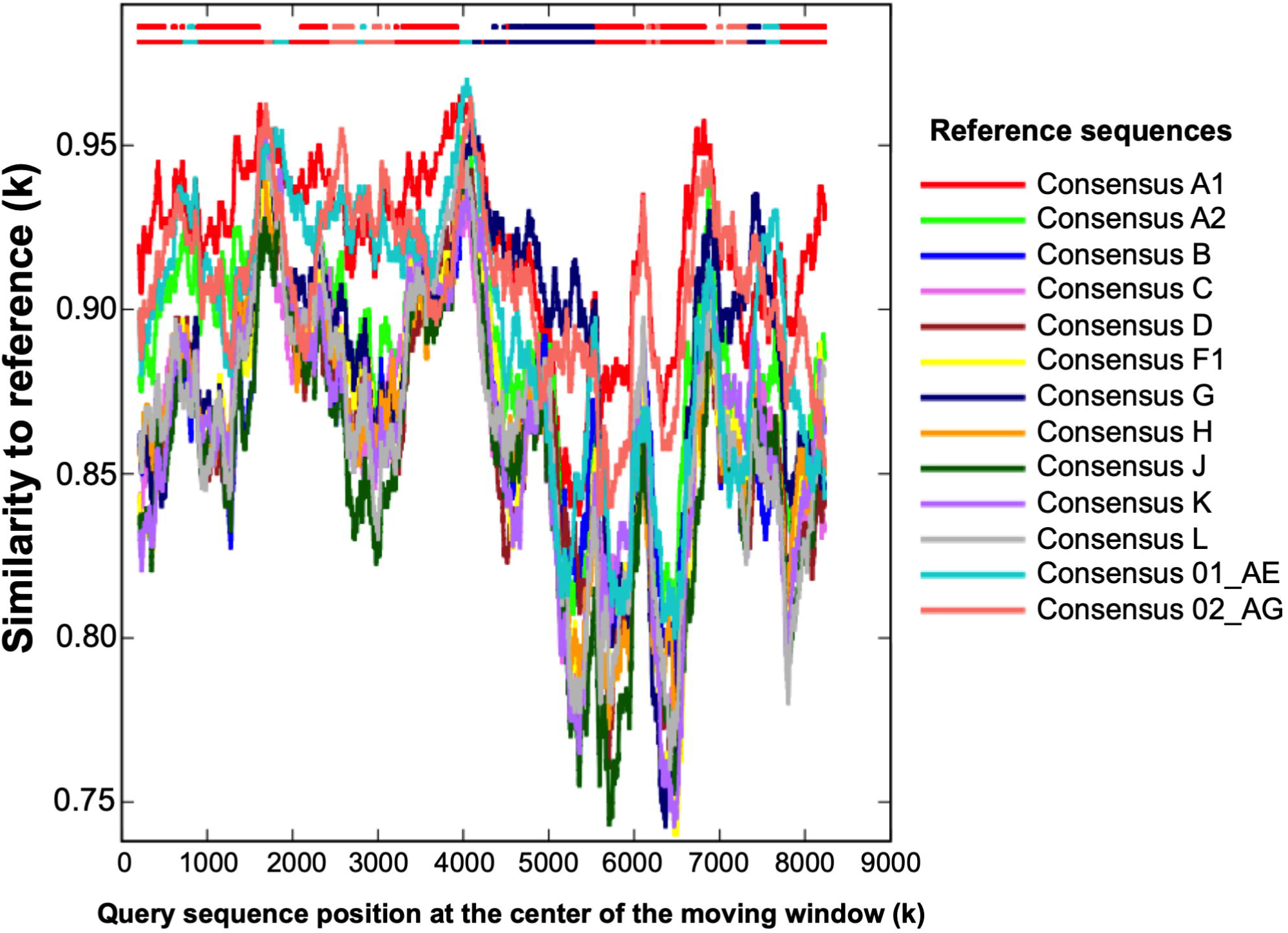
The participant’s HIV is a novel recombinant comprising subtypes A1, G, CRF01_AE, and CRF02_AG. Recombinant Identification Program (RIP) plot of the full genome sequence of a representative provirus isolated from the participant. The y-axis denotes the % similarity between the participant and each of the 13 reference sequences (each shown by a different color), using a sliding window of 1000 bases. The two lines at the top indicate the best matching reference sequence over a given region (lower bar) and whether this match meets the 90% confidence threshold (upper bar).

**Figure S2.**
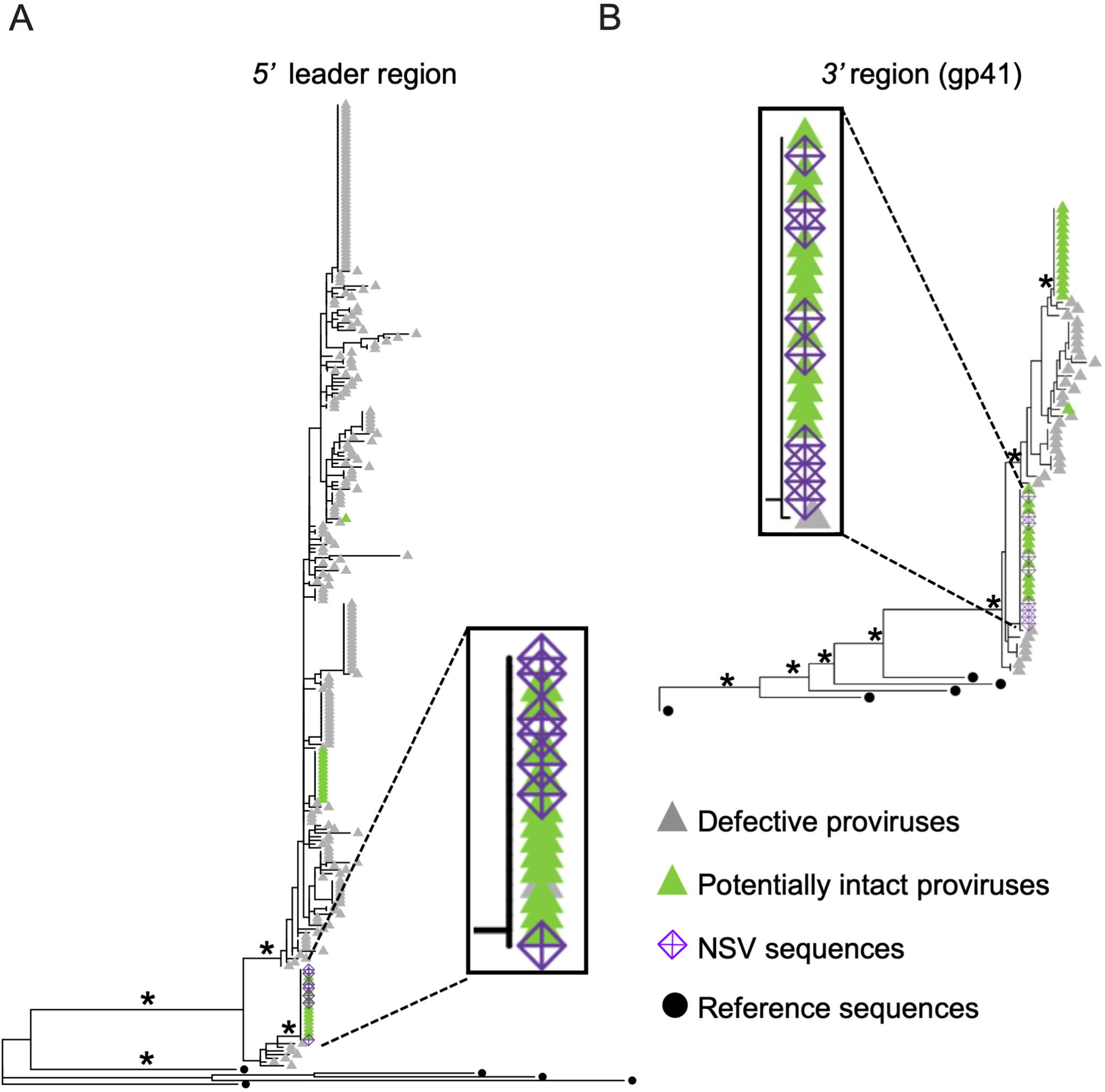
The plasma NSV sequence matches a clonally expanded provirus in partial 5’ leader and *env* regions. (A) Maximum likelihood within-host phylogeny inferred from a 545 base-pair region spanning partial 5’ leader and *gag* regions, in seven identical plasma sequences isolated during the NSV (purple crossed diamonds) and 255 non-hypermutated proviral sequences containing this region (green triangles for potentially intact proviruses; grey for defective ones). (B) Maximum likelihood within-host phylogeny inferred from a 498 base-pair region covering gp41, in nine identical plasma sequences isolated during the NSV and 61 non-hypermutated proviruses containing this region. (B). In both trees, black filled circles denote reference sequences and asterisks identify bootstrap values >80%. Scale in estimated substitutions per nucleotide site.

**Figure S3.**
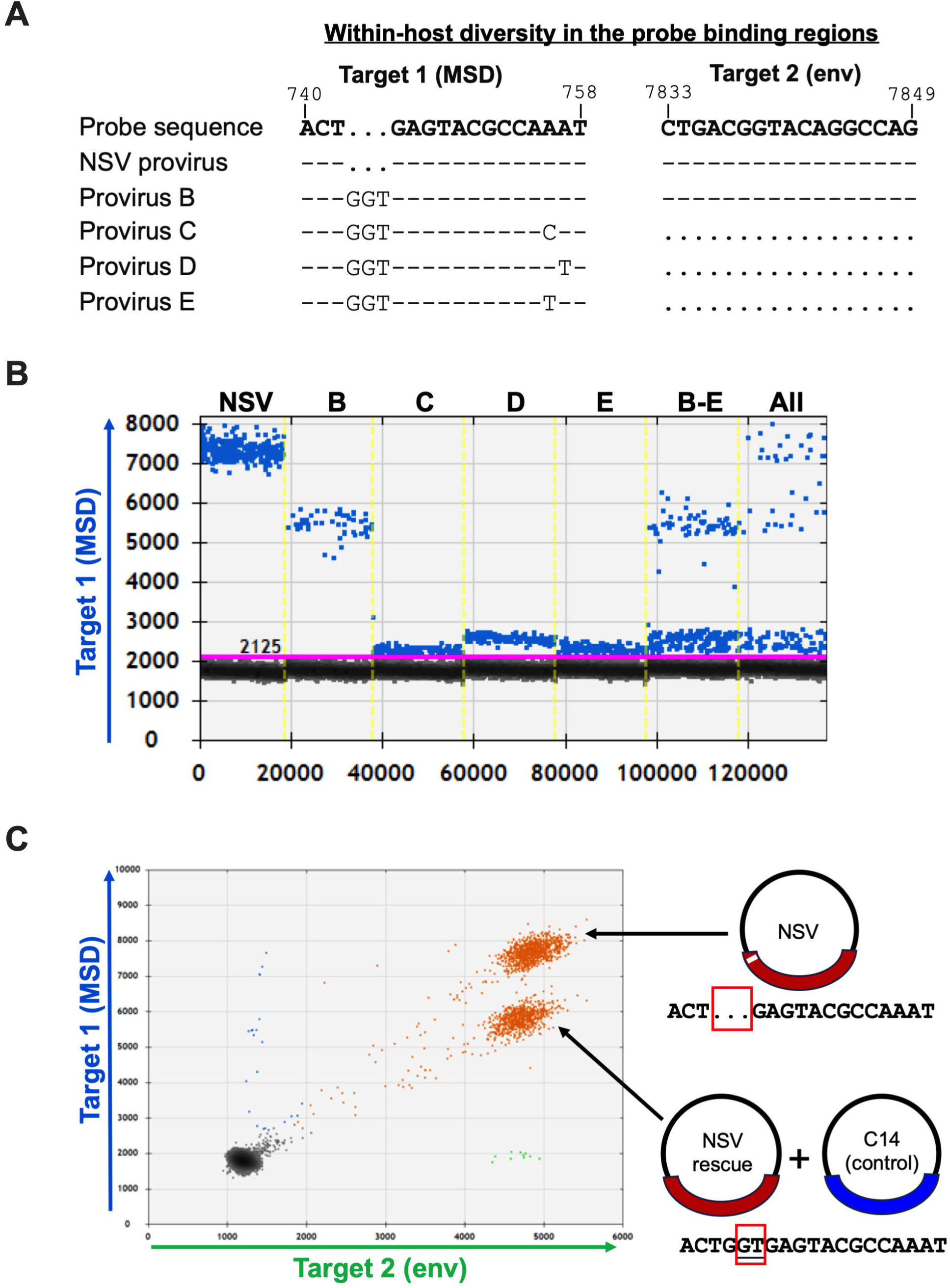
Validation of a custom ddPCR assay to detect the NSV provirus. Within-host sequence diversity in the assay’s MSD (target 1) and *env* (target 2) probe-binding regions, in the NSV provirus and the participant’s next four most abundant proviruses (labelled B-E). The target 1 (MSD) probe matched the NSV provirus exactly, while the target 2 (*env*) probe matched essentially all proviruses in this individual. (B) Representative one-dimensional ddPCR plots for target 1 (MSD) when the assay was applied to synthetic templates containing the sequences of the NSV provirus and proviruses B - E, either individually (first five columns) or when pooled (final columns labeled “B-E” and “All”). The NSV sequence consistently yielded signal amplitudes between 7000-8000 units, allowing it to be reliably discriminated from others even in a mixed pool. (C) Representative two-dimensional ddPCR plot when the assay was applied to a mixture of NSV, NSV rescue and C14 HIV molecular clones. Here, the plasmids are detected as double-positive (orange) events, where the NSV provirus is again consistently detected with signal amplitudes between 7000-8000 units for target 1. Single-positive (blue and green) events are due to occasional plasmid shearing.

**Figure S4.**
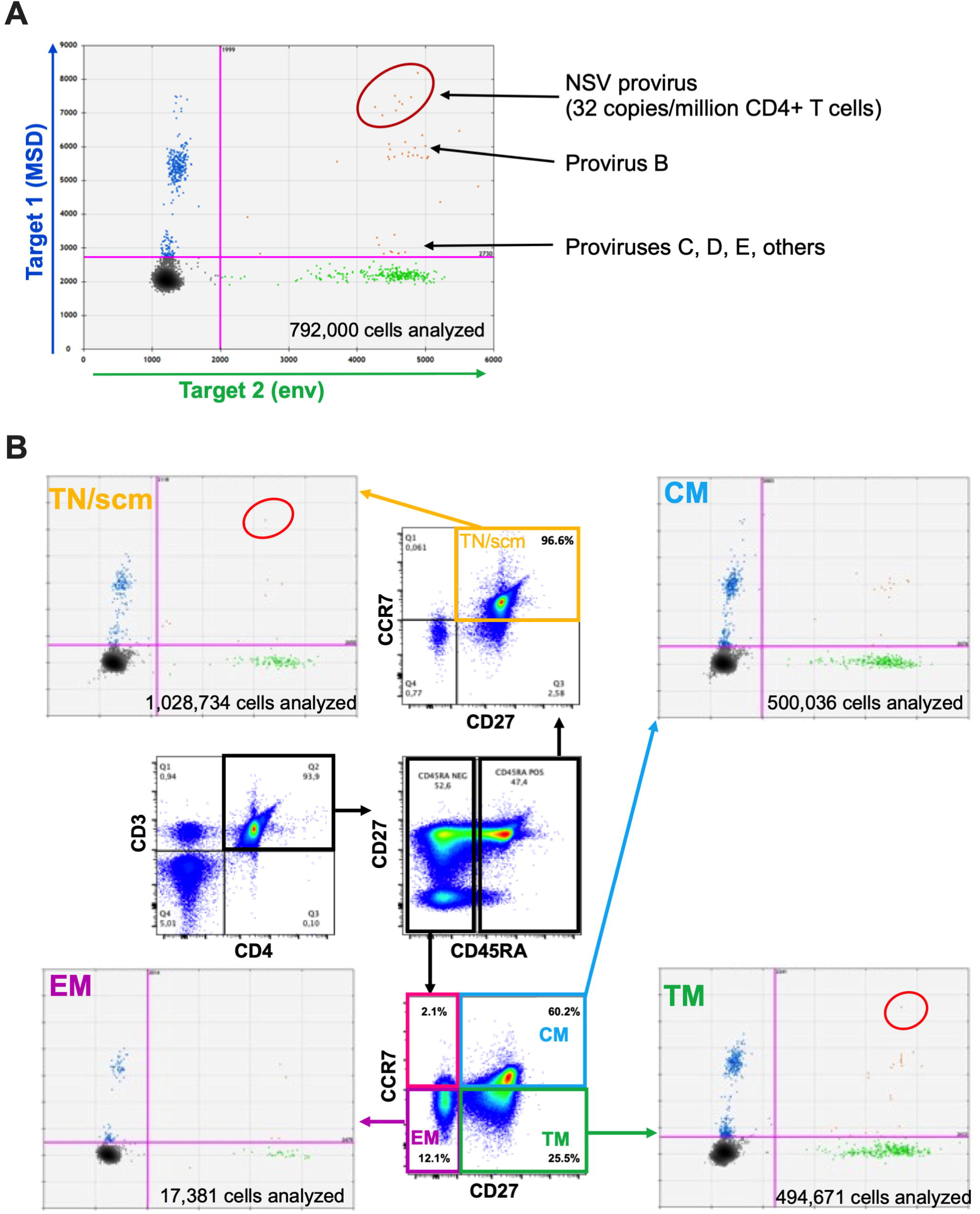
Applying a custom ddPCR assay to detect NSV provirus in isolated CD4+ T cells and subsets. (A) Representative raw ddPCR data from the analysis of bulk genomic DNA from 792,000 CD4+ T-cells, revealing that the NSV provirus was present at an estimated 32 copies per million total CD4+ T-cells. This is the same data as shown in Figure 5A. (B) Flow cytometry plots in the interior of the figure depict our initial sorting strategy for blood CD4+ T-cell subsets. Blood CD4+ T-cells were isolated by negative selection and then sorted into: Naïve or stem cell memory cells (TN/scm; CD45RA+CD27+CCR7+; boxed in yellow in the top flow plot); Central Memory cells (CM; CD45RA-CD27+CCR7+; boxed in blue in the bottom flow plot); Transitional Memory cells (TM; CD45RA-CD27+CCR7-+; boxed in green in the bottom flow plot) and Effector Memory cells (EM; CD45RA-CD27-CCR7-+; boxed in purple in the bottom flow plot). Matching colored arrows point to representative raw ddPCR data from each subsets, with the total number of cells analyzed per subset also shown. The pink box in the bottom flow plot identifies the atypical EM-like CD45RA-CD27-CCR7+ population that was not sorted in this experiment, but that we hypothesized could contain the NSV provirus clone population.

**Figure S5.**
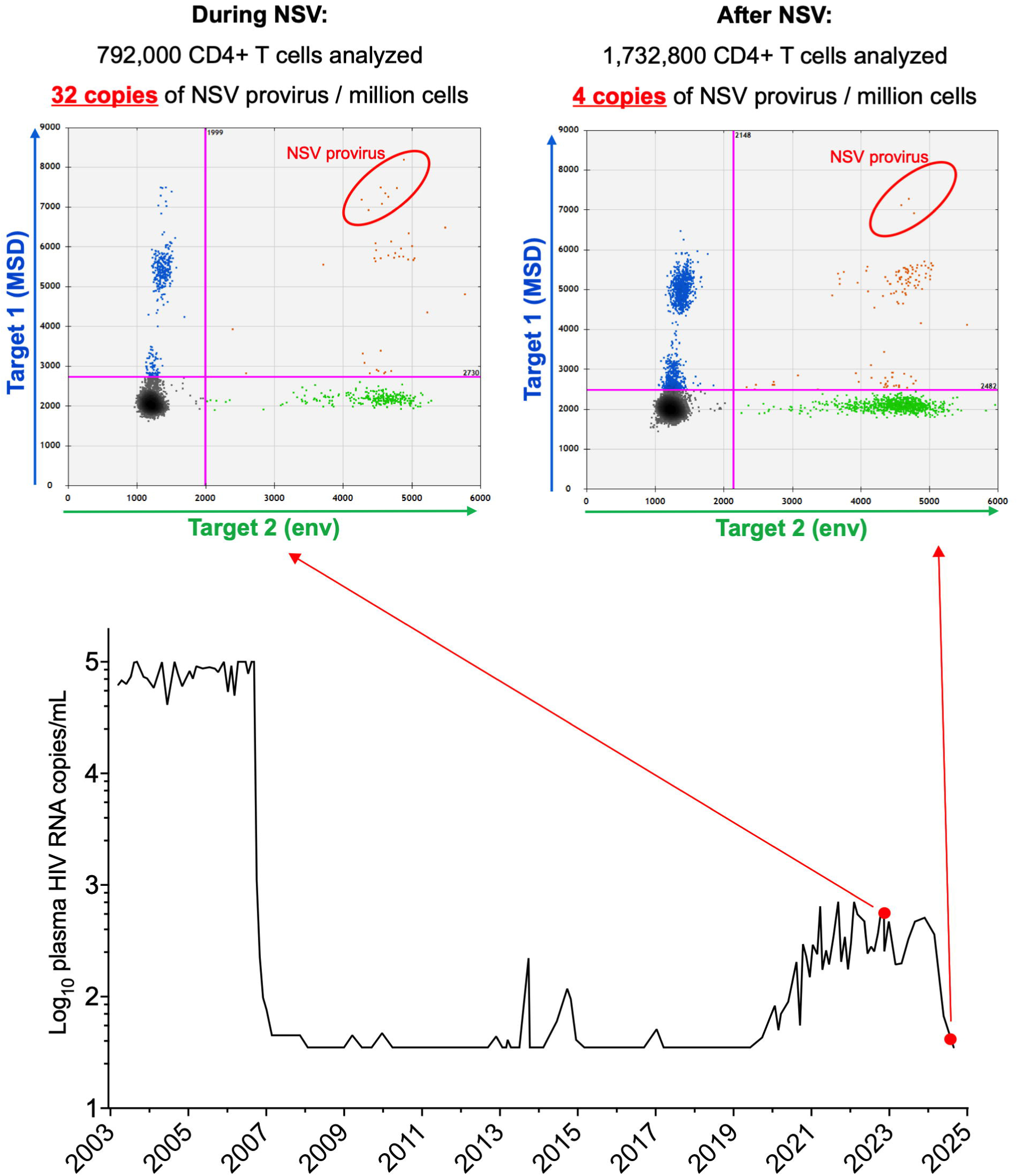
Spontaneous resolution of the non-suppressible viremia coincided with contraction of the NSV proviral clone. *Bottom:* The participant’s viral load history, with red circles indicating the two time points when blood CD4+ T-cells were analyzed for the presence of the NSV provirus. *Top left:* ddPCR plot showing NSV provirus frequency in total blood CD4+ T-cells during the viremic period (same data as in Figures 5A and S4A). *Top right:* ddPCR plot showing NSV provirus frequency in total blood CD4+ T-cells after the viremia had resolved.

**Figure S6.**
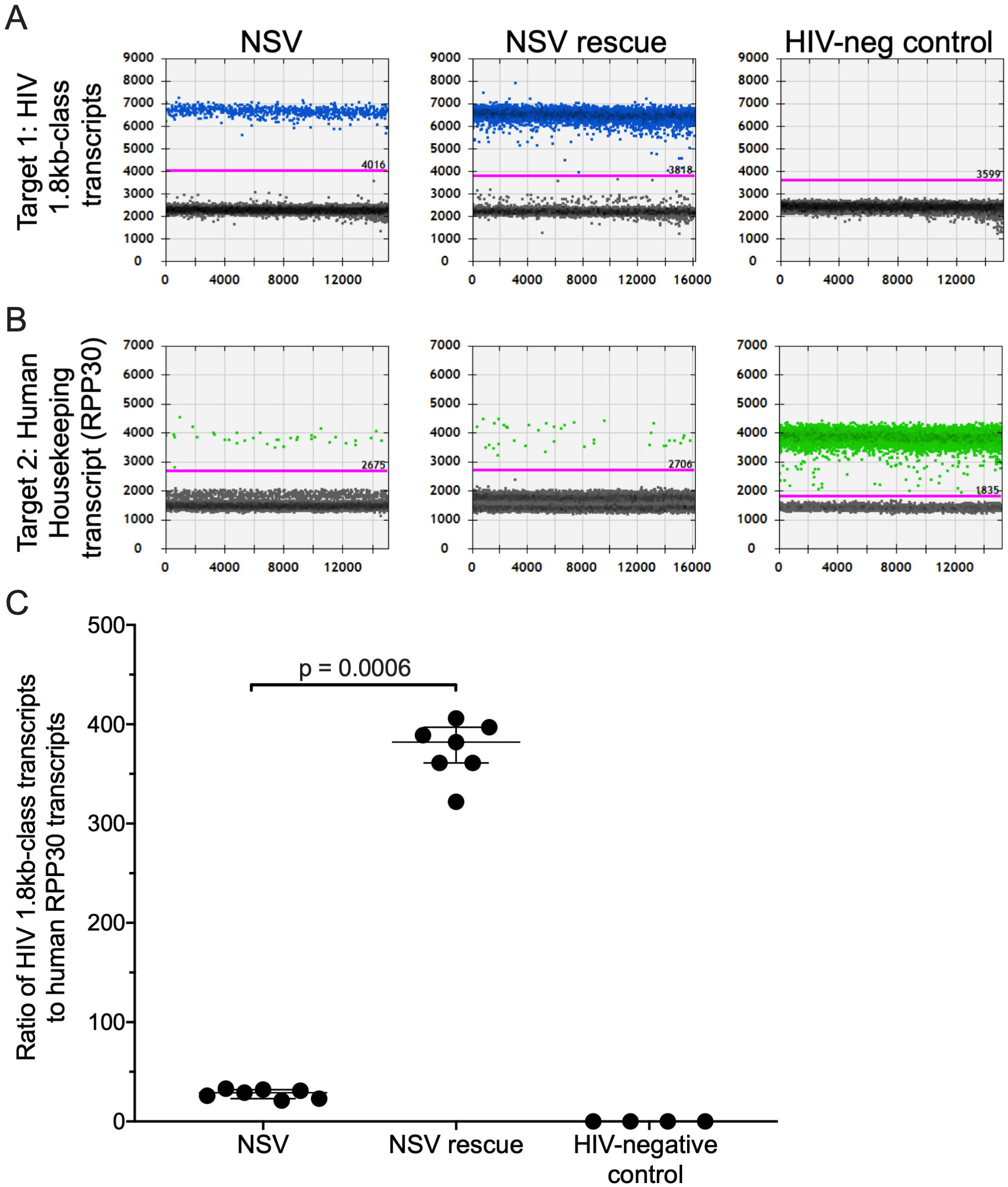
The MSD deletion substantially reduces spliced transcript abundance. (A) Representative ddPCR plots quantifying 1.8kb class HIV transcripts in HEK-293T cells transfected with equal amounts of NSV and NSV rescue plasmids, and in cells from an HIV-negative participant as a control. (B) Representative ddPCR plots quantifying transcripts for the human housekeeping gene RPP30 in these same cells. (C) Ratios of 1.8kb-class HIV transcripts to human RPP30 transcripts for the cell cultures shown in panels A and B, from a minimum of four (maximum six) technical replicates performed per cell type. Statistical significance was assessed using the Mann-Whitney U-test.

