## Supplemental Table S1 for "Prolonged non-suppressible viremia sustained by a clonally expanded, genomically defective provirus with an immune-evasive HIV protein expression profile"

**Table S1 - List of primers and probes**

| Assay | PCR reaction | Primer name | Direction | HXB2 coordinates | Sequence (5' to 3') |
|---|---|---|---|---|---|
| Near full-length amplification | Outer | 623-Fi(gag) | Forward | 623 - 629 | AAATCTCTAGCAGTGGCGCCCGAACAG |
| | | R9662-9686 | Reverse | 9686 - 9662 | TGAGGGATCTCTAGTTACCAGAGTC |
| | Nested | U5-638F | Forward | 638 - 666 | GCGCCCGAACAGGGACYTGAAARCGAAAG |
| | | U5-547R | Reverse | 9632 - 9604 | GCACTCAAGGCAAGCTTTATTGAGGCTTA |
| 5’ amplification | cDNA | GS3R | Reverse | 1841 - 1817 | TGACATGCTGTCATCATTTCTTCTA |
| | Outer | 623-Fi(gag) | Forward | 623 - 649 | AAATCTCTAGCAGTGGCGCCCGAACAG |
| | | GS3R | Reverse | 1841 - 1817 | TGACATGCTGTCATCATTTCTTCTA |
| | Nested | 638-666 | Forward | 638 - 666 | GCGCCCGAACAGGGACYTGAAARCGAAAG |
| | | GS1B | Reverse | 1339 - 1316 | AATCTTGTGGGGTGGCTCCTTCTG |
| Pol amplification | cDNA | RT3.1 | Reverse | 3859 - 3831 | GCTCCTACTATGGGTTCTTTCTCTAACTGG |
| | Outer | 5’CP1 | Forward | 1979 - 2005 | GAAGGGCACACAGCCAGAAATTGCAGGG |
| | | RT3.1 | Reverse | 3859 - 3831 | GCTCCTACTATGGGTTCTTTCTCTAACTGG |
| | Nested | 2.5 | Forward | 2011 - 2039 | CCTAGGAAAAAGGGCTGTTGGAAATGTGG |
| | | RT3798R | Reverse | 3798 - 3777 | CAAACTCCCACTCAGGAATCCA |
| 3’ amplification | cDNA | GP41RO | Reverse | 8819 - 8797 | CTTTTTGACCACTTGCCACCCAT |
| | Outer | GP41Fo | Forward | 7626 - 7648 | TTCAGACCTGGAGGAGGAGATAT |
| | | GP41RO | Reverse | 8819 - 8797 | CTTTTTGACCACTTGCCACCCAT |
| | Nested | GP41Fi | Forward | 7652 - 7674 | GGACAATTGGAGAAGTGAATTAT |
| | | GP41Ri | Reverse | 8771 - 8749 | CTGTCTTATTCTTCTAGGTATGT |
| LTR amplification | Outer | Auto FwdLTR | Forward | 1 - 27 | TGGATGGGTTAATTTACTCCARGAA |
| | | Auto Rev2Gag | Reverse | 1386 - 1367 | TTGCATAGCTGCCTGGTGTC |
| | Nested | Auto Fwd1LTR | Forward | 5 – 29 | TGGGTTAATTTACTCCARGAAAAGA |
| | | Auto RevGag | Reverse | 1317 - 1294 | TGATAGTGCTGTGAACATGGGTAT |
| HIV-1 spliced Transcripts: 4 kb class | cDNA | Auto-4REV-OUT | Reverse | 6101 - 6072 | CTACTAGTCCTACTATTGCAGATATAACTA |
| | Outer | 623-Fi(gag) | Forward | 623 - 649 | AAATCTCTAGCAGTGGCGCCCGAACAG |
| | | Auto -4REV-OUT | Reverse | 6101 - 6072 | CTACTAGTCCTACTATTGCAGATATAACTA |
| | Nested | U5-638F | Forward | 638 - 666 | GCGCCCGAACAGGGACYTGAAARCGAAAG |

| | | | | | |
|---|---|---|---|---|---|
| | | Auto -4REV-IN | Reverse | 6077 - 6046 | TAACTAAAGGTTGCATCACATATACTAATTAT |
| HIV-1 spliced Transcripts: 1.8 kb class | cDNA | Auto -1.8REV-OUT | Reverse | 5969 - 5947 | TGGTGATGGGTAAGGGTTGCTTT |
| | Outer | 623-Fi(gag) | Forward | 623 - 649 | AAATCTCTAGCAGTGGCGCCCGAACAG |
| | | Auto -1.8REV-OUT | Reverse | 5969 - 5947 | TGGTGATGGGTAAGGGTTGCTTT |
| | Nested | U5-638F | Forward | 638 - 666 | GCGCCCGAACAGGGACYTGAAARCGAAAG |
| | | Auto -1.8REV-IN | Reverse | 1668 - 1643 | TAAGGGTTGCTTTGGTACAGGATTTT |
| HIV-1 spliced Transcripts: 1 kb class | cDNA | EIR | Reverse | 9043 - 9015 | CCTTGTAAGTCATTGGTCTTAAAGGTACC |
| | Outer | 623-Fi(gag) | Forward | 623 - 649 | AAATCTCTAGCAGTGGCGCCCGAACAG |
| | | EIR | Reverse | 9043 - 9015 | CCTTGTAAGTCATTGGTCTTAAAGGTACC |
| | Nested | U5-638F | Forward | 638 - 666 | GCGCCCGAACAGGGACYTGAAARCGAAAG |
| | | GP41RO | Reverse | 8819 - 8797 | CTTTTTGACCACTTGCCACCCAT |
| NSV provirus detection (ddPCR) | RPP30 (human) | RPP30Fwd | Forward | NA | GATTTGGACCTGCGAGCG |
| | RPP30 (human) | RPP30Probe | NA (probe) | NA | VIC[a]-CTGACCTGAAGGCTCT-MGBNFQ[b] |
| | RPP30 (human) | RPP30Rev | Reverse | NA | GCGGCTGTCTCCACAAGT |
| | RPP30-Shear (human) | ShearFwd | Forward | NA | CCATTTGCTGCTCCTTGGG |
| | RPP30-Shear (human) | ShearProbe | NA (probe) | NA | FAM[c]-AAGGAGCAAGGTTCTATTGTAG-MGBNFQ |
| | RPP30-Shear (human) | ShearRev | Reverse | NA | CATGCAAAGGAGGAAGCCG |
| | HIV MSD | GagFwd | Forward | 692 - 711 | CAGGACTCGGCTTGCTGAAG |
| | HIV MSD | GagProbe | NA (probe) | 743 - 761 | FAM-ACTGAGTACGCCAAATTTT-MGBNFQ |
| | HIV MSD | GagRev | Reverse | 797 - 775 | GCACCCATCTCTCTCCTTCTAGC |
| | HIV env | EnvFwd | Forward | 7809 - 7825 | ACTATGGGCGCGGCGTC |
| | HIV env | EnvProbe | NA (probe) | 7849 - 7833 | VIC-CTGGCCTGTACCGTCAG-MGBNFQ |
| | HIV env | EnvRev | Reverse | 7939 - 7921 | CCCCAGACCGTGAGTTTCA |
| | HIV env | HypermutationProbe | NA (probe) | 7781 - 7798 | CCTTAGGTTCTTAGGAGC |
| HIV spliced transcript quantification (ddPCR) | HIV 1.8 kb spliced | Auto_MSFwd | Forward | 5978 - 5996 | AAGAAGCGGAGACAGCGAC |
| | HIV 1.8 kb spliced | Auto_MSProbe | NA (probe) | 6040 - 6045/ 8379 -8391[d] | AAAGCAACCCTTACCCATC |
| | HIV 1.8 kb | Auto_MSRev | Reverse | 8475 - 8456 | CGGTCACTAATCGAGTGGAT |

| | | | | |
|---|---|---|---|---|
| spliced | | | | |
| Human housekeeping RNA | RPP30Fwd | Forward | NA | GATTTGGACCTGCGAGCG |
| Human housekeeping RNA | RPP30Probe | NA (probe) | NA | VIC-CTGACCTGAAGGCTCT-MGBNFQ |
| Human housekeeping RNA | RPP30Rev | Reverse | NA | GCGGCTGTCTCCACAAGT |

[a]VIC = 2′-chloro-7′phenyl-1,4-dichloro-6-carboxy-fluorescein
[b]MGBNFQ = *3'* Minor groove binder; NFQ = Nonfluorescent quencher
[c]FAM = carboxyfluorescein
[d]Probe spans the D4/A7 junction of 1.8 kb multiply spliced transcripts
